# Directed growth during somatic cell fusion in *Neurospora crassa* requires contributions from two distinct MAP kinase pathways

**DOI:** 10.1101/2020.08.12.246843

**Authors:** Antonio Serrano, Hamzeh H. Hammadeh, Ulrike Brandt, André Fleißner

## Abstract

Somatic cell fusion in *Neurospora crassa* depends on two MAP kinases, MAK-1 and MAK-2, but their precise contributions to directed growth have remained unclear. Using analog-sensitive alleles and live-cell imaging, we found that MAK-1 activity is required for RAC-1 activation and polarized actin assembly, while MAK-2 controls the positioning of RAC-1 and actin at the fusion tips. Inhibition of MAK-1 disrupted cell communication, prevented actin focus formation, and blocked membrane recruitment of MAK-2 and SO. In contrast, MAK-2 inhibition caused SO and actin to shift to alternative cortical sites, showing that MAK-2 regulates their spatial orientation rather than activation. Actin disruption with latrunculin A eliminated MAK-2 and SO recruitment, confirming its essential role in their localization. Importantly, MAK-1 controlled actin polymerization only during tropic growth, not during general hyphal extension, and acted upstream of RAC-1. These findings establish complementary roles for MAK-1 and MAK-2 in directing RAC-1–dependent actin dynamics during fungal cell fusion.

**Summary statement:** MAK-1 activates the Rho GTPase RAC-1 to drive actin polymerization, while MAK-2 orients this RAC-1/actin complex toward the fusion partner, defining complementary MAPK roles in somatic cell fusion.

## Introduction

Cell polarization is a fundamental process in eukaryotic organisms, indispensable for growth, development, and certain types of cellular communication. It involves the asymmetric distribution of various molecular factors, including proteins, lipids, and RNA, to discrete regions of the cell cortex, enabling specialized functions at distinct cellular sites. A prominent example of this phenomenon is polarized growth, defined by selective cell expansion at define sites, resulting in elongated morphologies. Examples of cells undergoing highly polarized growth include neurons in animals [1], root hairs and pollen tubes formation in plants [2], and mating protrusions (shmoo) in yeast [3]. Filamentous fungi provide some of the most striking examples of polarized growth, as their hyphal cells extend continuously at the tip and form branched, interconnected mycelial networks capable of rapid and directional exploration of the environment [4–6].

Polarized growth in fungi can be broadly divided into two categories: general polarized growth and directed growth (chemotropic or tropic growth). During general polarized growth, hyphae or the germ tubes of germinating spores protrude continuously in a defined axis, often without changing direction. By contrast, directed growth refers to the reorientation of hyphal tips in response to specific environmental or cellular cues, such as host-derived signals in pathogenic fungi, nutrient gradients, or the presence of a compatible fusion partner. This facilitates cell-cell fusion (somatic fusion) of compatible cells, a pivotal process for colony networking, resource sharing, and the formation of specialized sexual structures [7].

In filamentous fungi, this process of somatic fusion takes place repeatedly throughout development and serves as the basis for building an interconnected colony. Fusion occurs first between germinating spores (germling fusion) and later between mature hyphae (hyphal fusion), establishing cytoplasmic continuity between cells and allowing the exchange of nutrients, signals, and organelles within the mycelial network. Each fusion event follows a defined sequence that includes mutual recognition between compatible partners, chemotropic growth of the cell tips toward one another, and subsequent cell wall remodeling and membrane merger. The recognition and tropic growth phases are driven by a “cellular dialogue,” a dynamic oscillatory communication in which interacting cells alternate between signal-sending and signal-receiving states to coordinate directed growth and ensure successful fusion [8, 9].

A key aspect of understanding how cells translate extracellular cues into polarization is the mitogen-activated protein (MAP) kinase network [10, 11]. This conserved signaling machinery governs multiple facets of polarized growth across diverse organisms. In the baker’s yeast *Saccharomyces cerevisiae*, the Fus3 MAPK pathway is the most studied regulator of mating, wherein cells detect peptide pheromones released by opposite mating type. Upon pheromone sensing by specific G protein-coupled receptors, a cascade of GTPases and kinases converges on phosphorylation of Fus3p, driving transcriptional changes and orchestrating the reorganization of the actin cytoskeleton at the site of the pheromone gradient. Concurrently, the cell wall integrity (CWI) pathway, centered on the MAPK Slt2/Mpk1, collaborates with the pheromone pathway to ensure proper cell wall remodeling [12] and the control of polarity under mechanical stress [13]. Additionally, the CWI pathway regulates cell wall modifications required for the culmination of the sexual cell fusion process. Tight coordination of both MAPK pathways ensures that membrane extension and cell wall restructuring take place correctly, enabling precise polarize growth toward the mating partner [14]. Related mechanisms governing MAPK-dependent polarization operate in other yeast and fungal species. In the fission yeast *Schizosaccharomyces pombe*, MAPK such as Pmk1 (the CWI pathway homolog to yeast) contribute to cell integrity and polar growth [15]. Similarly, the filamentous fungus *Neurospora crassa* relies on two MAPK modules, the MAK-1 (Mpk1p homolog) and MAK-2 (Fus3p homolog) pathways, for hyphal fusion and colony morphogenesis. Mutants lacking either pathway fail to initiate normal somatic fusion events and cannot form the interconnected mycelial networks of wild-type colonies [16, 17].

Although all fungal hyphae exhibit continuous tip extension (general polarized growth), certain stimuli necessitate a shift to directed growth. In *N. crassa*, vegetative germling fusion requires tropic interactions between germ tube tips. The switch to directed growth involves rearranging the polarity machinery, including classic polarity regulators, such as Rho-type GTPases, including CDC-42 and RAC-1, that govern actin cytoskeleton organization and exocytosis, ensuring the delivery of secretory vesicles to the cell tips [18, 19]. Notably, in this fungus, CDC-42 is essential for sustaining general germ tube extension, while RAC-1 drives the tropic response and directed growth associated with cell fusion events [19]. Both regulators seem to interact with similar downstream components, such as the formin Bni1, that nucleates actin polymerization, and associated filaments form actin cables that enable secretory vesicles delivery to the polarity sites [19].

During this process, the MAPK MAK-2 oscillates between the cytoplasm and the plasma membrane at the tip of the interacting cells, facilitating rapid switches in the cellular dialogue [8, 9]. Meanwhile, the MAPK MAK-1 pathway functions predominantly at later fusion stages and contact recognition, controlling the cell wall remodeling and membrane merger once the fusion partners come into physical contact [20]. However, it is currently unclear, if the MAK-1 pathway also plays any role during directed growth of the fusion cells towards each other.

By delineating temporal and functional roles of these MAPK pathways, we provided a more comprehensive framework for understanding the cellular transition between general and directed polarized growth. By employing a chemical genetics approach with analog-sensitive MAPK variants (*Shokat* alleles) [21, 22], in combination with high spatial resolution live-cell imaging, we demonstrate that MAK-2 is decisive in orienting growth toward the fusion interacting cell, ensuring that RAC-1 (Rho-type GTPase) remain properly polarized to drive precise chemotropic growth. Conversely, MAK-1 is also required for maintaining the polarized state during directed growth and ensuring actin oscillation between the interacting cells. Taken together, our results support a model in which the two MAPK act in tandem during directed polarized growth.

## Results

### Inhibition of MAK-1 kinase activity disrupts the process of cell fusion

The CWI MAPK MAK-1 is known to play important roles in the successful completion of cell fusion, including an essential but unknown role in establishing cell fusion competence, and its activity is crucial for growth arrest of fusing cells after physical contact is established [20, 23]. However, it remains unclear whether MAK-1 also plays a role during the tropic interaction of fusion partners. Deletion of the *mak-1* gene completely disrupts cell fusion competence [23], prompting us to create an analog-sensitive version, or *Shokat* allele [21], of MAK-1 to manipulate its activity in a spatial-temporal manner (Fig. S1A and B).

The resulting strain, MAK-1^E104G^ (*mak-1::mak-1^E104G^-hph*), appeared macroscopically wild type-like, indicating that the mutated kinase is functional in the absence of the inhibitor (ATP-analog, 1-NM-PP1) and supplemented with 1% DMSO (the solvent of 1-NM-PP1) as a control (Fig. S1C). In contrast, addition of the inhibitor (final concentration of 40 µM) to the growth medium resulted in a Δ*mak-1*-like phenotype, whereas the wild-type control strain remained unaffected (Fig. S1C). Microscopic analysis showed that the presence of the inhibitor drastically reduced the cell fusion rate of the MAK-1^E104G^ strain, enabling us to analyze the functions of MAK-1 at specific stages of the cell fusion process, such as the cell-cell communication phase (Fig. S1D). Additionally, as previously shown in other publications, neither the solvent DMSO nor the ATP-analog had any effect on wild-type cells, demonstrating the specificity of the inhibitor for analog-sensitive variants in our fungal system.

To investigate further the effects of MAK-1 inhibition on cell-cell communication, multiple strains were created. The proteins MAK-2 and SO are crucial to the cell fusion process, and thus, we examined the localization of these following MAK-1 inhibition. The strains MAK-1^E104G^ + SO-GFP and MAK-1^E104G^ + MAK-2-GFP were created through sexual crosses (see Materials and methods for more details). All strains exhibited comparable MAK-1 inhibition macroscopically phenotypes upon the addition of the ATP-analog (data not shown). When live-cell imaging of cell-cell communication, the addition of DMSO did not affect the normal oscillatory recruitment of MAK-2 (Fig. 1A). However, the addition of the ATP-analog completely disrupted the cell-cell communication process, and the dynamic localization of MAK-2 was interrupted. The kinase was no longer observed on the membrane, and no full detachment occurred even after almost 18 minutes (Fig. 1B). Similarly, the addition of DMSO did not affect the usual oscillatory recruitment of SO (Fig. 1C). However, 10 minutes after the addition of the kinase inhibitor, SO accumulated at the membrane in both cell tips, and the GFP signals were more widely distributed around the cell tips (Fig. 1D). These findings suggest that MAK-1^E104G^ activity is essential for proper MAK-2 and SO dynamics during the directed growth phase. Similarly, the characteristic MAK-2/SO dynamics also occur during hyphal fusion [24]. We therefore examined the effects of MAK-1 inhibition on hyphal fusion events. When DMSO was added, MAK-2 oscillated in a wild-type manner (Fig. S2A). However, when MAK-1^E104G^ was inhibited, MAK-2 exhibited a disruption of their dynamics similar to the findings in germlings (Fig S2B). Regarding SO, DMSO did not affect the usual oscillatory dynamics (Fig. S2C), but the addition of the inhibitor completely disrupted the process (Fig. S2D). While MAK-2 completely vanished from the hyphal tip, SO remained as single dots distributed widely around the plasma membrane of both interacting hyphae. These data indicate that the function played by MAK-1 during germling fusion is also conserved in hyphal fusion.

**Figure 1.**
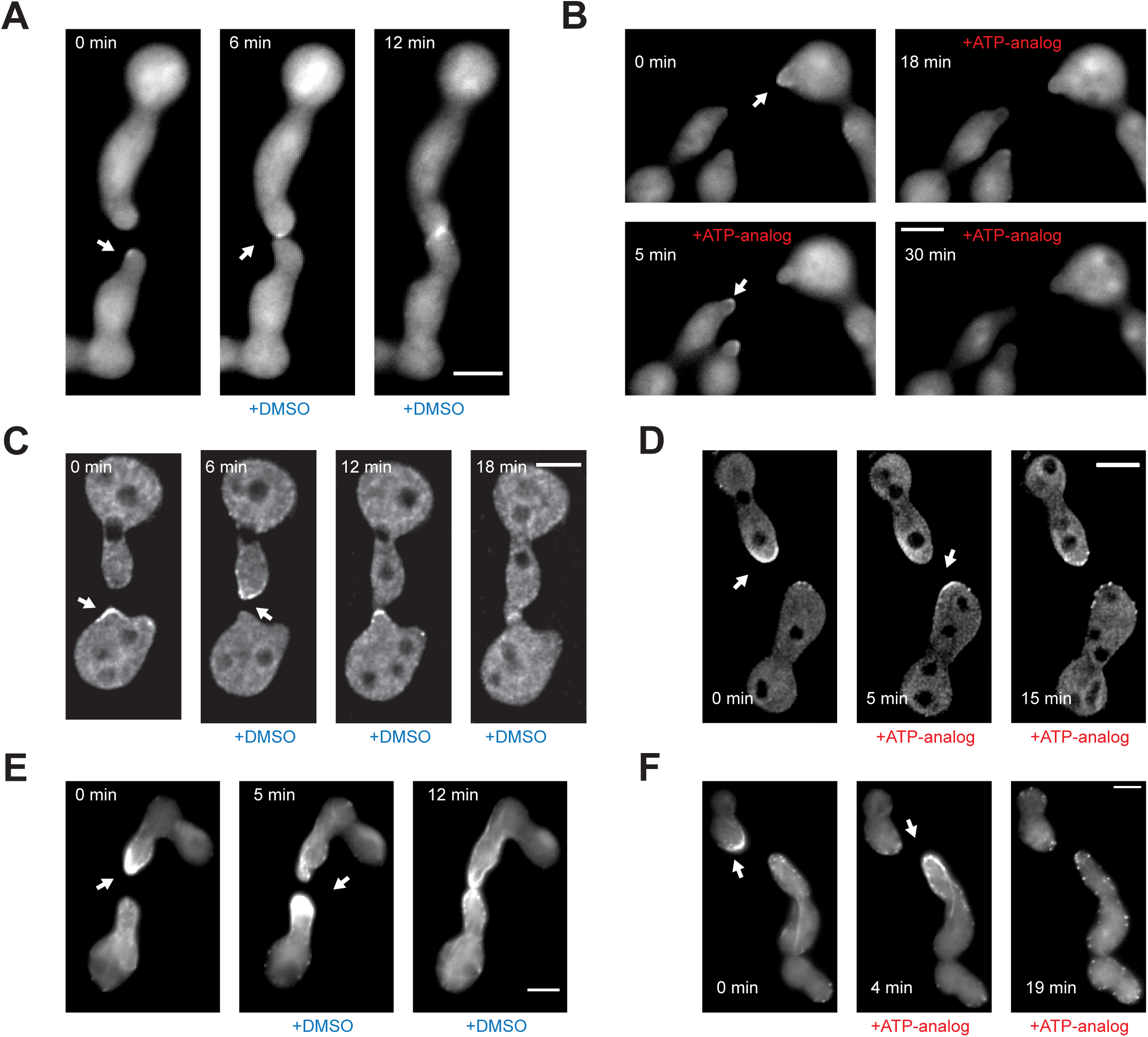
Activity inhibition of MAK-1 disrupts the oscillatory localization of MAK-2/SO and LifeAct. **(A)** Oscillatory localization of MAK-2-GFP during cell fusion in a MAK-1^E104G^background (strain 865) with the addition of DMSO. **(B)** Inhibition of MAK-1^E104G^ disrupts the oscillatory dynamic localization of MAK-2. **(C)** Oscillatory dynamic localization of SO during cell fusion in a MAK-1^E104G^ background (strain 874) with the addition of DMSO. **(D)** Inhibition of MAK-1^E104G^ disrupts the oscillatory dynamic localization of SO. **(E)** Oscillatory localization of LifeAct-GFP during cell fusion in a MAK-1^E104G^ background (strain 869) with the addition of DMSO. **(F)** Inhibition of MAK-1^E104G^ disrupts the oscillatory dynamic localization of LifeAct-GFP. All observations were made multiple times (n≥15). Scale bars 5 µm.

Proper actin organization is crucial for polarized hyphal extension in fungi. In non-interacting cells, actin is primarily found at the tips of growing germ tubes in the form of actin patches localized around the cell tips and actin cables [18]. During cell fusion, the disruption of actin assembly leads to a failure of polarized, directed growth of the fusion tips, while microtubules are dispensable for this process [25]. To image actin dynamics, we used the LifeAct reporter, previously adapted for our fungal system [18], and created a strain MAK-1^E104G^ + LifeAct-GFP through a sexual cross (see Materials and methods for more details). This strain behaved similarly to the wild type in the presence of DMSO, and typical actin cables were present at the cell tips during the communication phase and during cell merger, comparable to wild-type cells. Previously unreported in our system, we observed that actin dynamics showed a comparable oscillatory and alternating accumulation at the cell tips of the interacting cells as MAK-2 and SO (Fig. 1E). When MAK-1^E104G^ was inhibited, actin accumulation was clearly affected: the accumulation of actin cables from the tips disappeared and actin patches were no longer focused on the cell tips but localized in a wider distribution around the cell membrane of the entire cell body (Fig. 1F). Actin oscillation was also observed in hyphal fusion, and inhibition of MAK-1^E104G^ disrupted actin accumulation at the hyphal tips, in a similar fashion as described in germlings (Fig. S2E and F).

These data indicate that MAK-1 is involved in the regulation of the actin cytoskeleton, MAK-2 and SO dynamics during the cell fusion process.

### Actin accumulation at the cell tips oscillates out-of-phase with SO/MAK-2 dynamics

Remarkably, the observation of actin localization in the presence of DMSO in the MAK-1^E104G^ inhibitable strain suggested that actin accumulates at the cell tips of the interacting cells and that this accumulation oscillates in an antiphase manner (Fig. 1E). To further investigate whether this was a specific effect of this strain, we examined actin localization in wild-type cells. Using the wild-type strain LifeAct-GFP [18], we made the same observations (Fig. 2A & S3A). Based on these data, we hypothesized that actin accumulation likely occurs together with SO, since the presence of SO at the cell tips is associated with a signal sending state of the fusing cell [26, 27]. We therefore created a heterokaryon combining LifeAct-GFP, and dsRed-SO, to enable observation of actin and SO within the same cell. When interacting cell pairs were observed, dynamic interferes co-localization was found, in which SO was recruited to the membrane at the tip of the first pair, while actin was recruited in the second interacting cell. After a few minutes, both proteins co-localized at the tip of the first partner. Finally, actin disassembled from the membrane of the upper cell and was recruited in the lower cell, while SO remained at the tip of the upper cell (Fig. 2B and S3B, C and D). This indicates that actin polymerization at the cell tips is out-of-phase with the usual dynamics of MAK-2/SO. To test whether actin oscillates during vegetative growth, germinated spores of wild-type LifeAct-GFP were observed under the fluorescence microscope, in single cells with no interacting partner nearby, and no oscillation was reported (Fig. 2C), indicating that oscillatory accumulation of actin is exclusive to cell fusion growth.

**Figure 2.**
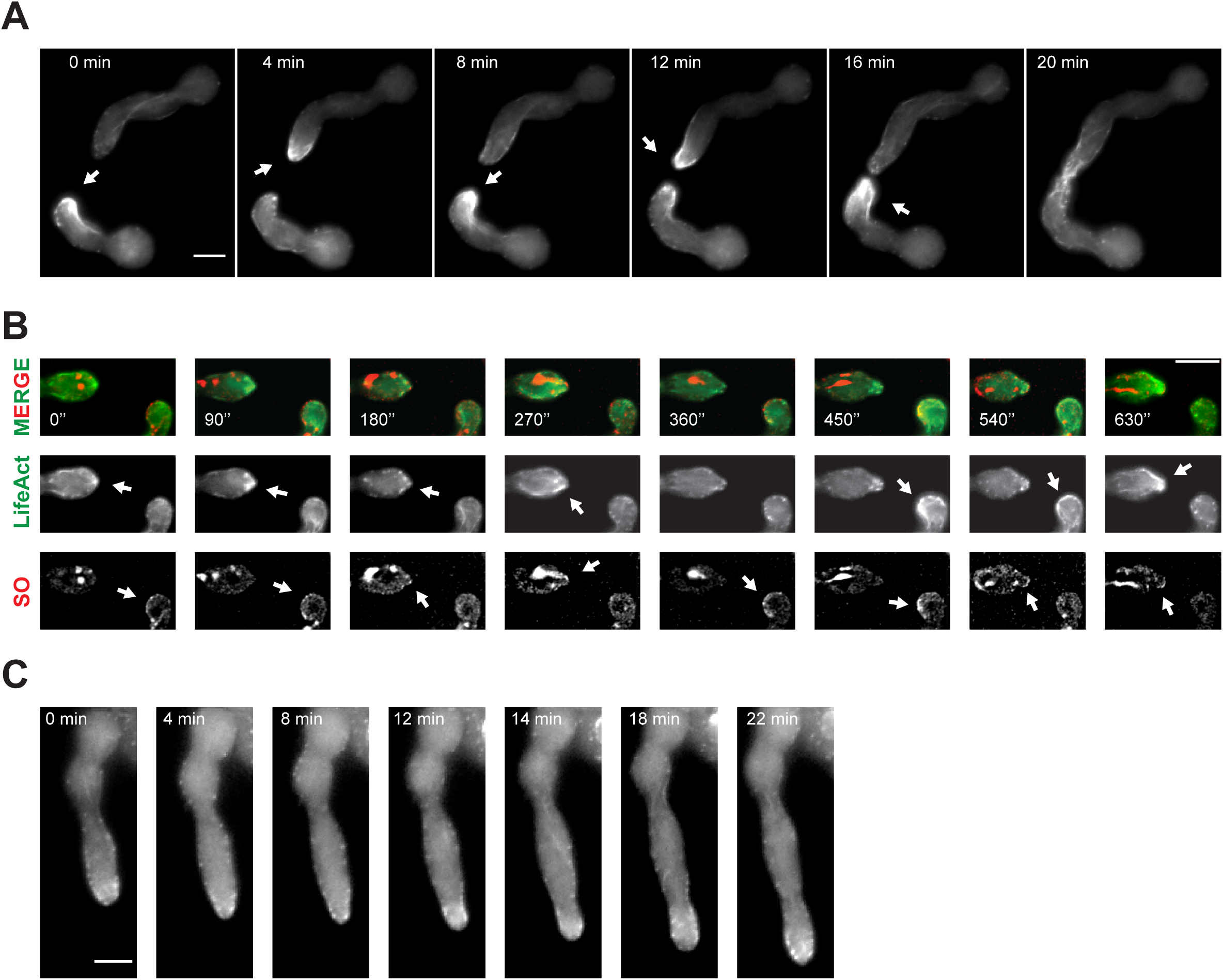
Actin focalizes to the tips of the interactings cells in an oscillatory manner. **(A)** Oscillatory dynamic localization of actin fusion aster at the cell tips of the interacting cells with LifeAct-GFP (strain 754). Similar observations were made multiple times (n=10) **(B)** Lifeact-GFP co-localization with dsRed-SO during the interaction of fusing cells in the heterokaryon created with strains 754 and N1-30. **(C)** LifeAct-GFP localization in a non-interacting cell during vegetative growth (strain 754). Similar observations were made multiple times (n=10). Scale bars 5 µm

### Inhibition of MAK-2 disrupts the cell fusion process

Based on previous results, it was hypothesized that MAK-1 may directly participate in actin polymerization or that actin misregulation may be an indirect effect of the disruption of the cell fusion process caused by MAK-1 inhibition. To test this hypothesis, a similar strategy was followed by creating an analog-sensitive version of MAK-2 to decipher SO and actin dynamics during the tropic growth phase. The generated MAK-2^Q100A^ (*Pmak-2-mak-2::Ptef-1-mak-2^Q100A^-hph*) strains exhibited a macroscopic phenotype similar to wild type in the presence of DMSO and a Δ*mak-2*-like phenotype when the kinase was inhibited (Fig. S4A-C). Microscopically, cell fusion rates were not fully comparable to the wild-type strain in the presence of DMSO (73.8% vs 51.3%; WT/MAK-2^Q100A^), but cell fusion was still observed. In contrast, the cell fusion rate was comparable to Δ*mak-2* when the kinase inhibitor was added (Fig. S4D). These data indicate that the strain is functional for cell fusion in the absence of the kinase inhibitor and allow us to analyze the cell dynamics when MAK-2^Q100A^ is inhibited in a spatial-temporal manner.

To evaluate the impact of MAK-2^Q100A^ inhibition on the subcellular localization of SO and actin, we generated strains MAK-2^Q100A^ + SO-GFP and MAK-2^Q100A^ + LifeAct-GFP. We observed that in the presence of DMSO, the dynamic localization of SO was similar to the wild-type phenotype (Fig. 3A). However, upon addition of the kinase inhibitor, the oscillation of SO rapidly ceased (within 4 minutes), and in 10 out of 15 observations, the localization of SO remained broadly distributed around the cell membrane (Fig. S5A). Interestingly, in 5 out of 15 observations, the localization of SO shifted to a different area of the cell, suggesting that the complex formed at the cell tips may travel together around the cell membrane after MAK-2 inhibition (Fig. 3B). We also examined the localization of actin and observed fusing cell pairs under the microscope in the presence of DMSO, displaying the previously described oscillatory accumulation of actin at the cell tips (Fig. 3C). However, in the presence of the kinase inhibitor, the actin localization shifted to a different position within the cell, with 10 out of 20 observations displaying actin travelling to the opposite cell tip in a manner similar to that of SO (Fig. 3B). After 10 minutes, actin still accumulated at the new position, despite the termination of tropic interaction and directed growth (Fig. 3D). In some cases (10 out of 20), actin shifted to a different position but remained close to the cell tip (Fig. S5B). These data suggest that MAK-2 inhibition has a slightly different impact on the disruption of actin localization and indicate that MAK-1 and MAK-2 may have distinct roles in cytoskeleton organization during cell fusion. Moreover, the observed effect on actin localization rules out the hypothesis of an indirect effect on actin cytoskeleton due to the disruption of cell fusion.

**Figure 3.**
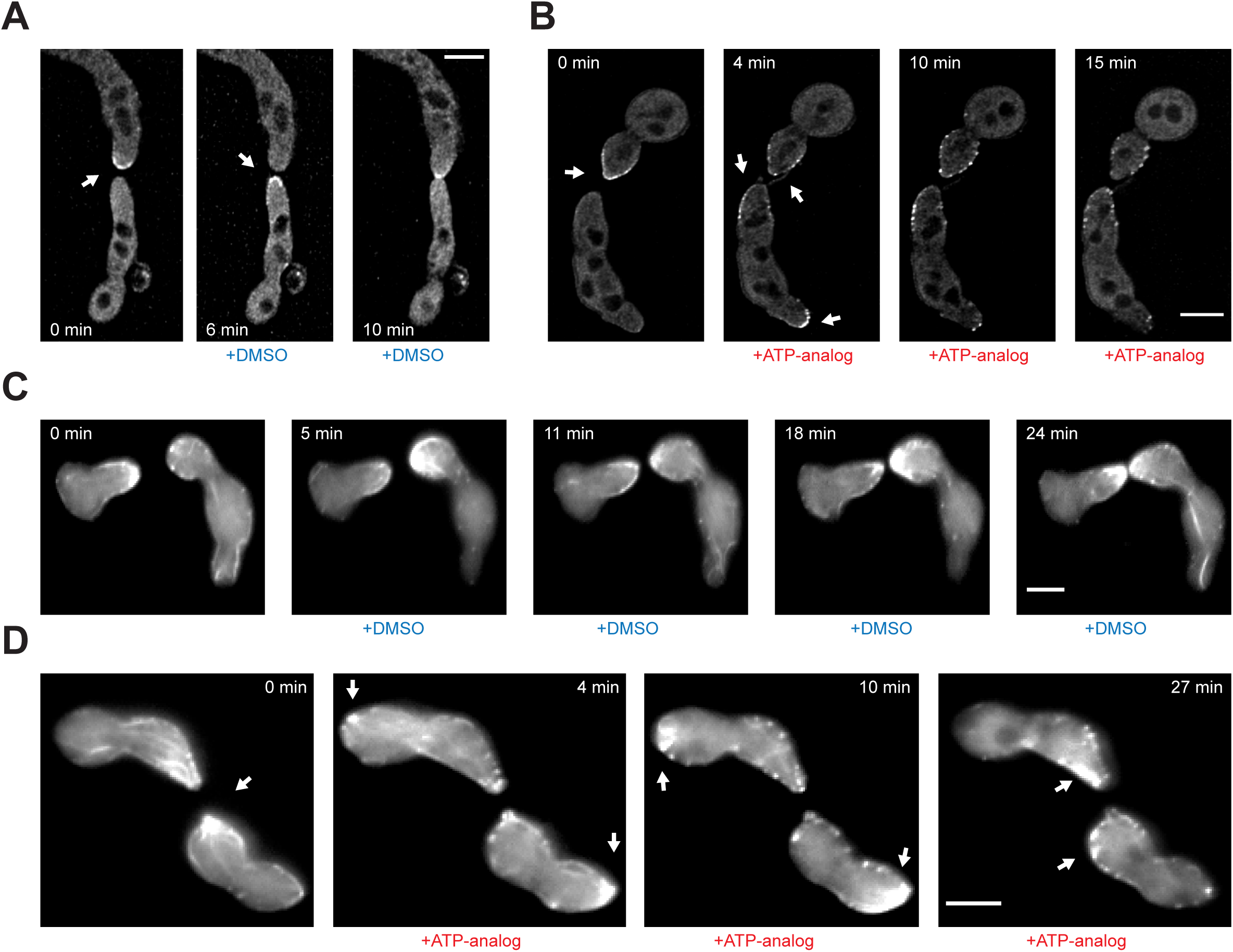
Activity inhibition of MAK-2 disrupts the SO dynamic localization and mislocalizes actin around the cell membrane. (A) Oscillatory localization of SO-GFP during cell fusion in a MAK-2^Q100A^ background (strain 911) with the addition of DMSO. (B) Inhibition of MAK-2^Q100A^ disrupts the oscillatory dynamic localization of SO-GFP. (C) Oscillatory localization of LifeAct-GFP during cell fusion in a MAK-2^Q100A^ background (strain 915) with the addition of DMSO. (D) Inhibition of MAK-2^Q100A^ disrupts the oscillatory dynamic localization of LifeAct-GFP. All observations were made multiple times (n≥10). Scale bars 5 µm.

### The actin cytoskeleton plays a crucial role in the dynamic localization of MAK-2 and SO

The inhibition of MAK-1^E104G^ disrupts proper membrane recruitment of MAK-2/SO, while also disturbing actin accumulation at the cell tips. Actin cables serve as tracks for the transport of multiple cargoes, including potential secretory vesicles that may be involved in the cell fusion process [18, 28]. Our data support two alternative hypotheses: either MAK-1 activity is directly involved in both MAK-2/SO recruitment and actin organization, or it is only involved in actin organization, and actin disruption results in disturbed MAK-2/SO localization.

In yeast, Fus3p recruitment is mainly mediated through actin cables organized at the Shmoo of the mating cells [29]. To test whether actin is required for proper MAK-2/SO recruitment, we employed the actin-disrupting drug latrunculin A [30]. In a pretest, we determined that a concentration of 10 μM resulted in the dissociation of F-actin in germinating spores within 6-10 minutes (Fig. S5C). When we tested MAK-2 localization, we observed that actin disruption due to latrunculin A resulted in the disassembly of MAK-2 after 8 minutes, and no further MAK-2 membrane recruitment was observed in the cells (Fig. 4A). Similarly, the dynamic localization and membrane recruitment of SO were also affected by actin inhibition (Fig. 4B), suggesting that proper actin polymerization at the cell tips is essential for membrane recruitment of MAK-2 and SO. To further understand the implications of MAK-1/MAK-2 in this process, we decided to investigate the roles of the upstream regulators of actin polymerization, the Rho-GTPases.

**Figure 4.**
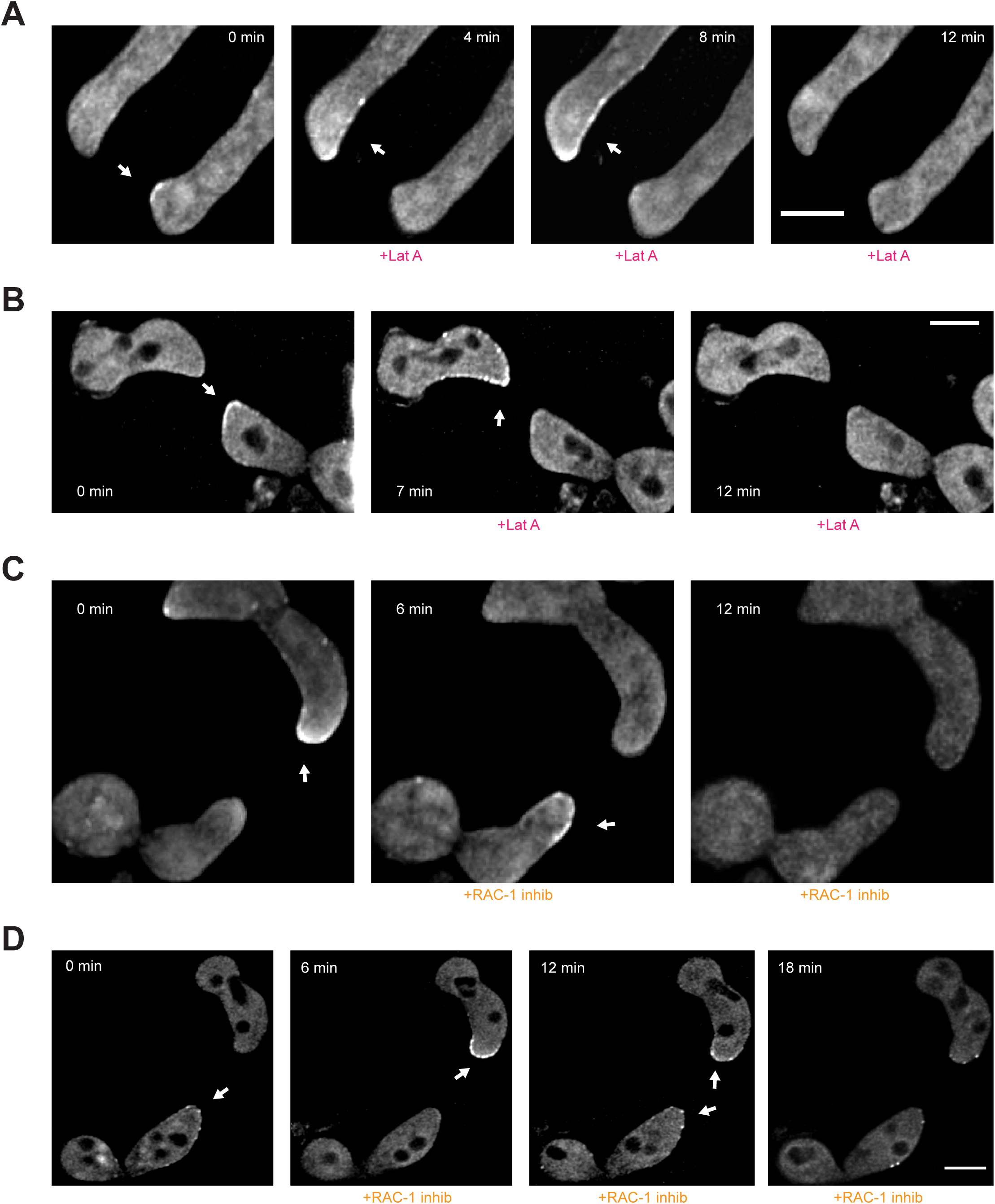
Inhibition of actin assembly disrupts the MAK-2/SO dynamic localization. **(A)**Inhibition of actin with latrunculin A (Lat A) disrupts the localization of MAK-2-GFP (strain 865) **(B)** Inhibition of actin with Lat A disrupts the localization of SO-GFP (strain 874). **(C)** Inhibition of RAC-1 with RAC-1 inhibitor (NSC23766) disrupts the localization of MAK-2-GFP (strain 865) **(D)** Inhibition of RAC-1 with RAC-1 inhibitor disrupts the localization of SO-GFP (strain 874). All observations were made multiple times (n≥10). Scale bars 5 µm.

Rho GTPases are conserved eukaryotic proteins that regulate actin dynamics by cycling between active (GTP-bound) and inactive (GDP-bound) states [3]. Upon activation, they translocate to the plasma membrane and interact with actin regulators, such as formins, to promote actin cable formation. In filamentous fungi, RAC-1 and CDC-42 localize to growing tips but play distinct roles in cell fusion and germ tube growth, respectively [19]. The RAC-1–specific inhibitor NSC3766 has been used to distinguish their functions. In our experiments, we tested if specific inhibition of RAC-1 with NSC3766 affects the dynamics of actin and the recruitment of MAK-2, SO, and actin assembly. In interacting cell pairs expressing MAK-2-GFP, the protein was recruited to the membrane in a wild-type manner before addition of RAC-1 inhibitor. The addition of the RAC-1 inhibitor at a concentration of 100 μM resulted in defects in the recruitment of MAK-2 after 12 minutes. MAK-2 fully disappeared from the membranes of both cells and the interaction ended (Fig. 4C). When SO dynamics were tested, the usual membrane accumulation at the cell tips was observed. Like the previous experiment, defects in the localization of SO were observed 12 minutes after the addition of the RAC-1 inhibitor. However, in contrast to MAK-2, SO remained widely distributed at the membranes of both cells although the interaction had ended (Fig. 4D). To note, this mislocalization of SO was comparable to the one observed after MAK-1 inhibition (actin patches distributed all around the cell membrane), suggesting that both proteins (MAK-1 & RAC-1) may play functions within the same pathway. We also examined actin localization in the presence of the RAC-1 inhibitor. The addition of the inhibitor resulted in a decline in the number of actin cables, and actin patches seems to be distributed more widely around the plasma membrane. After 12 minutes, actin cables had fully disappeared and directed cell growth was terminated (Fig. S5D).

These data indicate that the proper polymerization of actin at the cell tips of the interacting cells, through activation of the Rho GTPase RAC-1, is essential for the dynamic localization of MAK-2 and SO.

### MAK-1 activity is required for RAC-1 activation, whereas MAK-2 activity determines its positioning

Previously, we described a potential connection between the activity of MAK-1 and RAC-1. Both proteins are necessary for actin assembly during cell fusion, and similar defects are observed when each one is inhibited. In contrast, MAK-2 inhibition disrupts the cell communication process, although the actin complex becomes unstable within the cell. Together, these data support two potential hypotheses: MAK-1 and RAC-1 function in the same pathway, either upstream or downstream, controlling actin polymerization, or RAC-1 and MAK-1 regulate, independently, a common target that is essential for actin assembly, such as the formin BNI-1. To test these hypotheses, we decided to analyze the dynamics of active RAC-1 after MAK-1 and MAK-2 inhibition using the Cdc42-Rac-interacting-binding (CRIB) reporter. If the first hypothesis is correct, MAK-1 inhibition will inactivate RAC-1, causing reporter detachment from the membrane. Since Rho GTPases associate with membranes only when active, detachment indicates inactivation. Alternatively, if the second hypothesis is correct, MAK-1 inhibition will not affect RAC-1, and the reporter will remain at the plasma membrane, suggesting that MAK-1 and RAC-1 act independently on a common factor involved in actin polymerization during cell fusion.

To note CRIB reporters in filamentous fungi binds as a GEF protein to activated Rho-GTPases without disrupting their function, thereby allowing the localization of both activated RAC-1 and CDC-42 at the same time in our fungal system [19]. The use of the highly specific RAC-1 inhibitor allowed to distinguish the functions of both proteins, since CDC-42 is unaltered by this chemical. As a control, we first tested the already reported effects of RAC-1 inhibition on the CRIB reporter. As previously described, the addition of the RAC-1 inhibitor to interacting cells resulted in the disappearance of the reporter from the apical tip and cell growth arrest (Fig. S6A). The complete disappearance of the reporter from the tips of the interacting cells indicates that only RAC-1, and not CDC-42, is present at the growth cone of the fusing cells [19].

In our study, we observed that the CRIB reporter was present permanently in both cell tips during fusion, as previously shown [19] (Fig. 5A). Interestingly, following inhibition of MAK-1, the CRIB reporter disappeared from the plasma membrane and cell growth was arrested (Fig. 5B). It is noteworthy that both RAC-1 and MAK-1 inhibition produced a similar effect on the CRIB reporters, suggesting that MAK-1 inhibition specifically affects RAC-1 activity, leading to the disruption of the actin assembly, membrane recruitment of MAK-2 and SO, and the general fusion process. Consistent with our hypothesis, we predict that MAK-2 inhibition does not affect the activity of RAC-1 but rather its localization, as previously observed with the actin in earlier experiments. As previously observed, the localization of the CRIB reporter remained permanently at both cell tips during the entire tropic growth phase in our control (Fig. 5C). However, when MAK-2 was inhibited, the CRIB reporter remained recruited to the plasma membrane, and its localization appeared to be widely distributed around the cell tip (Fig. 5D). These suggest that MAK-2 may control the localization and, therefore, growth directionality of RAC-1.

**Figure 5.**
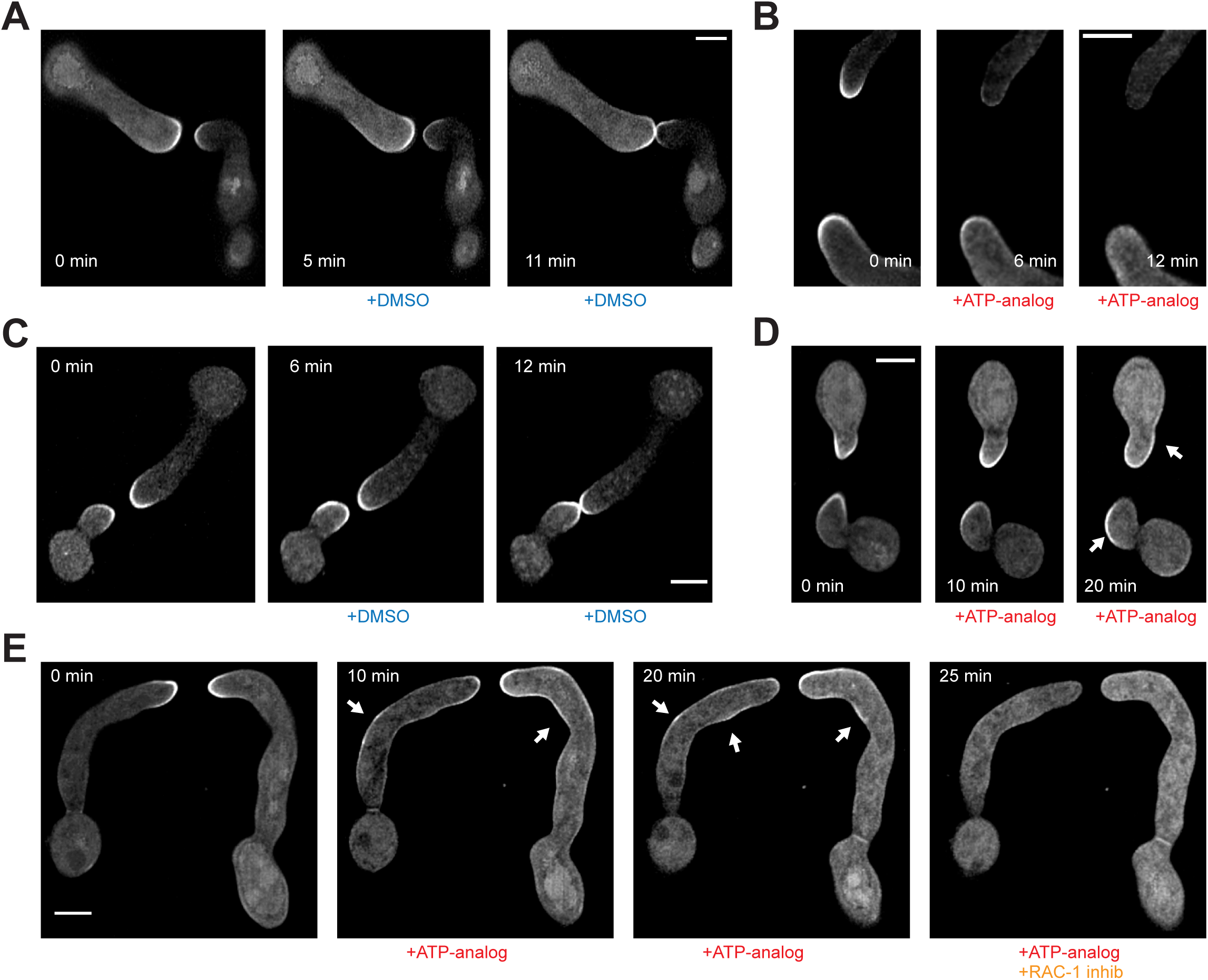
Activity inhibition of MAK-1 results in RAC-1 inactivation while MAK-2 inhibition results in RAC-1 mislocalization. **(A)** Localization of the CRIB-GFP reporter during cell fusion in a MAK-1^E104G^ background (strain 943) with the addition of DMSO. **(B)** Inhibition of MAK-1^E104G^ disrupts CRIB-GFP localization during cell fusion. **(C)** Localization of the CRIB-GFP reporter during cell fusion in a MAK-2^Q100A^ background (strain 949) with the addition of DMSO. **(D)** Inhibition of MAK-2^Q100A^results in mislocalization of the CRIB-GFP during cell fusion. **(E)** Inhibition of MAK-2^Q100A^ results in mislocalization of the CRIB-GFP during cell fusion, and subsequent addition of the RAC-1 inhibitor results in disruption of the CRIB reporter. Comparable observations were made multiple times (n≥10). Scale bars 5 µm.

Nevertheless, as we have previously mentioned, the CRIB reporter allows for the localization of the activated forms of both RAC-1 and CDC-42 at the same time. To determine if RAC-1, and not CDC-42, changes its position after MAK-2 inhibition, we applied the RAC-1 inhibitor after inhibiting MAK-2, combining both chemicals in a temporal resolution. Fusing cells were observed under the microscope, and the kinase inhibitor was added. After observing the switching in the position of the CRIB reporter provoked by MAK-2 inhibition, the RAC-1 inhibitor was added. Interestingly, the CRIB reporter disappeared completely from the plasma membrane, indicating that MAK-2 inhibition of fusing cells affects exclusively the position of RAC-1 at the plasma membrane (Fig. 5E & S6B). These data indicate that MAK-2 inhibition in tropically growing cells specifically affects the localization of RAC-1, which remains active and recruited to the plasma membrane but loses its growth directionality towards the cell fusion partner and changes its position to another area in the cell.

Taken together, all these data support a model in which the inhibition of MAK-1 specifically affects RAC-1 membrane recruitment, which results in the disassembly of the actin focus organized by the Rho-GTPase and the arrest of growth of the interacting cells.

### MAK-1 is a specific regulator of actin polymerization during cell fusion

We have shown that MAK-1 activity is crucial for proper actin polymerization at the tips of interacting cells during cell fusion. This leads to the question of whether MAK-1 functions specifically in directed growth or whether it plays a role in general polar growth. To address this question, we conducted an experiment to observe the effect of kinase and RAC-1 inhibitors (as a negative control) independently on non-interacting cells. The length of the germ tube was quantified under a microscope before and after incubation with the kinase inhibitor, RAC-1 inhibitor, or DMSO. To ensure that only non-interacting cells were tested, we analyzed germlings that were at a distance of more than 30 μm from the nearest cell. Our results indicate that there were no significant differences observed between the DMSO, kinase inhibitor, or RAC-1 inhibitor treatments, supporting the notion that MAK-1/RAC-1 exclusively regulates actin polymerization and therefore tropic growth during cell fusion (Fig. 6A). Additionally, we conducted a similar experiment to test whether MAK-2 had any effect on growth rate and obtained similar results (Fig. 6A).

**Figure 6.**
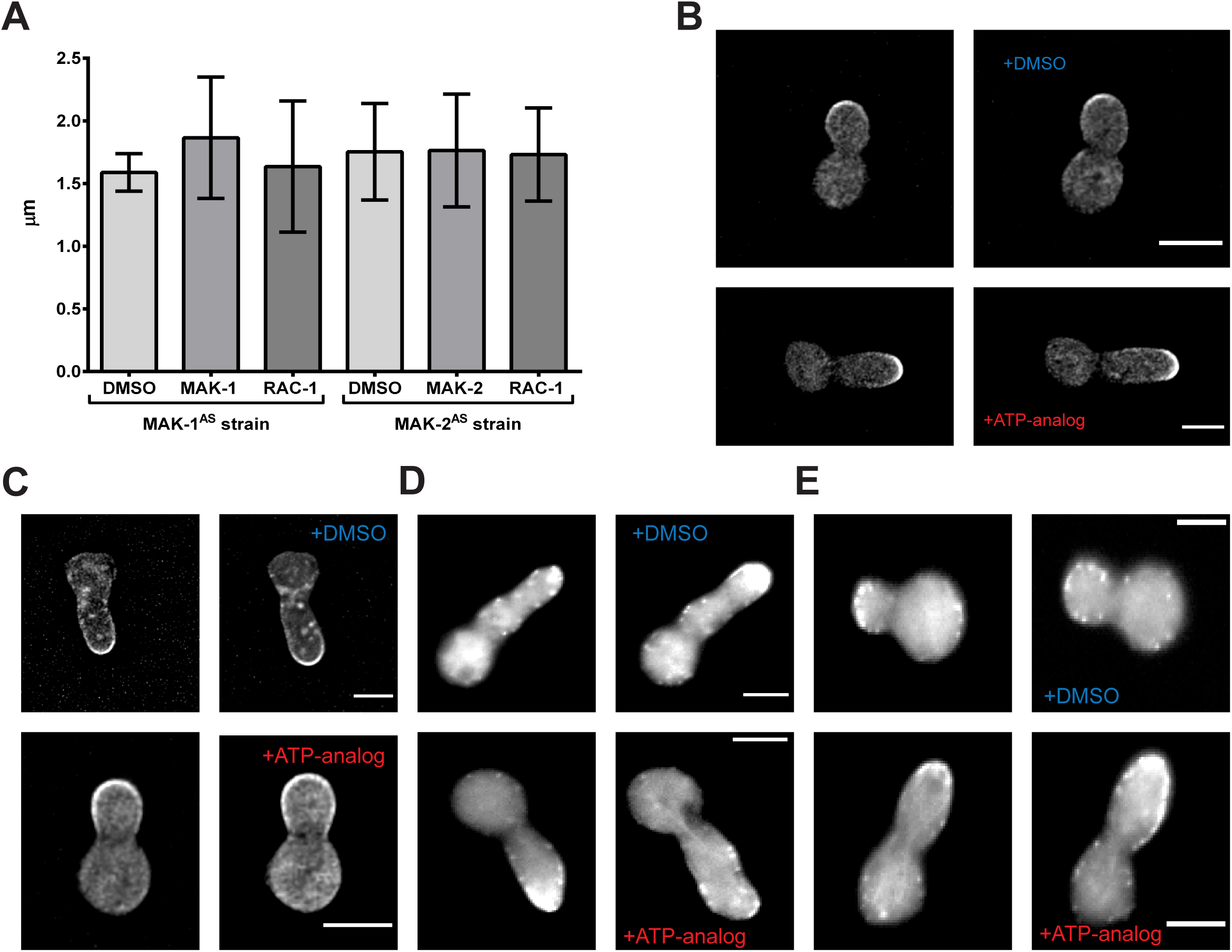
Inhibition of either MAK-1 or RAC-1 does not affect actin localization in vegetative growth. **(A)** Quantification of cell elongation in cells treated either with DMSO, ATP-analog (MAPK inhibitor) or RAC-1 inhibitor. No significant differences are observed between all tests (n≥25). **(B)** Upper panel: localization of the CRIB-GFP reporter in non-interacting cells before and after addition of DMSO in a MAK-1^E104G^ background (strain 943). Bottom panel: localization of the CRIB-GFP reporter in cells before and after inhibition of MAK-1^E104G^. **(C)** Upper panel: localization of the CRIB-GFP reporter in non-interacting cells before and after addition of DMSO in a MAK-2^Q100A^background (strain 949). Bottom panel: localization of the CRIB-GFP reporter in cells before and after inhibition of MAK-2^Q100A^**(D)** Upper panel: localization of LifeAct-GFP in non-interacting cells before and after addition of DMSO in a MAK-1^E104G^background (strain 869). Bottom panel: localization of LifeAct-GFP in cells before and after inhibition of MAK-1^E104G^. **(E)** Upper panel: localization of LifeAct-GFP in non-interacting cells before and after addition of DMSO in a MAK-2^Q100A^ background (strain 915). Bottom panel: localization of LifeAct-GFP in cells before and after inhibition of MAK-2^Q100A^. Similar observations were made multiple times (n≥5). Scale bars 5 µm.

Furthermore, we investigated the localization of the CRIB reporter and actin before and after treatment with DMSO, kinase inhibitor, or RAC-1 inhibitor, in non-interacting cells. As expected, there were no consistent differences observed in the localization of the CRIB reporter when cells were treated with DMSO or the kinase inhibitor (Fig. 6B) and with H_2_O (the solvent for the RAC-1 inhibitor) or the RAC-1 inhibitor (Fig. 6C) in the MAK-1-inhibitable strains. Similarly, no significant differences were observed in the localization of the CRIB reporter when the cells were incubated with DMSO or the kinase inhibitor (Fig. 6B), H_2_O, or the RAC-1 inhibitor (Fig. 6C) in the MAK-1^E104G^ and MAK-2^Q100A^ (Fig. 6D and E).

To further confirm these findings from the chemical genetics approach, we studied actin organization in the *mak-1* knockout mutant obtained from the *N. crassa* gene knockout collection (NCU09482). A Δ*mak-1* LifeAct-GFP strain was created through sexual crosses (see Materials and methods for more details). As previously reported, this mutant exhibited a cell fusion-deficient phenotype [23]. The spores of this isolate were incubated in MM for 2 hours, and actin organization was studied using fluorescence microscopy. In the Δ*mak-1* mutant cells, germ tube elongation and actin assembly at the growing cell tips were comparable to wild type [18] (Fig. S6C).

Taken together, these data indicate that MAK-1 exclusively functions as an actin polymerization regulator during the cell fusion process and is not involved in actin polymerization during general vegetative growth. In other words, MAK-1 is not involved in the same actin-regulatory pathway as CDC-42.

### MAK-1 functions upstream of RAC-1

MAK-1 has been identified as a key factor in regulating the communication and growth arrest phases during cell fusion. This kinase plays a crucial role in the growth arrest phase, which involves remodeling of the cell wall to allow for the completion of the fusion process. Analog-sensitive versions of MAK-1 have been shown to partially inhibit the growth arrest phase, resulting in disrupted cell merger and the emergence of a corkscrew-like phenotype in fusing cells, also known as twisting germ tubes [20].

To investigate whether MAK-1 and RAC-1 are part of the same pathway, we conducted experiments using wild-type cells and partial inhibition of RAC-1. We tested three different concentrations of RAC-1 inhibitor (100, 50, and 25 µM), with only the latter concentration allowing for the induction of cell fusion events (Fig. 7A). Furthermore, a large number of cells exhibited defects in polar growth, which may be related to the effect of RAC-1 inhibition. To further investigate this observation, we quantified the percentage of cells that showed tropic defects (observed as growth-forming circles) after RAC-1 inhibition in medium with Ca^2+^. Approximately 12.5% of cells exhibited this growth, compared to the water control. To determine whether this defect was caused by inhibition of the general germ tube elongation machinery, we performed similar quantifications on wild-type cells placed on media without Ca^2+^ (which inhibits the induction of cell fusion) and on a cell fusion mutant, such as Δ*so*. We observed that polarity defects were not observed in either case, suggesting that this phenotype may be caused by partial inhibition of RAC-1 specifically when cells are undergoing cell fusion (Fig. 7B). Although MAK-1 and RAC-1 function within the same actin-regulatory pathway during cell fusion, there is currently a lack of strong evidence to determine which protein is upstream/downstream. Interestingly, both proteins are localized at the cell tips during the cell growth arrest phase [19, 20]. Therefore, we conducted experiments to investigate how the localization of MAK-1 and RAC-1 were affected after inhibition of RAC-1 or MAK-1, when the cells are merging. To observe the effects of the subcellular localization of MAK-1/RAC-1, cells that had just established contact were observed under a microscope and either RAC-1 or the kinase inhibitor was added. When RAC-1 was observed, its localization remained at the contact area of the fusing cells that had just made contact, and the addition of DMSO had no effect on its localization (Fig. 8A). However, the complete inhibition of MAK-1 in fusing cells rapidly induced mislocalization of RAC-1, which completely disappeared from the plasma membrane while cells likely stopped their cellular interaction (Fig. 8B). The addition of water, the solvent of the RAC-1 inhibitor, did not affect the cell fusion event and MAK-1 localization during the final stages of cell fusion (Fig. 8C). Interestingly, the addition of the RAC-1 inhibitor in interacting cells did not have any significant effect on the localization of MAK-1, which remained at the contact area, although cells likely stopped their interaction (Fig. 8D).

**Figure 7.**
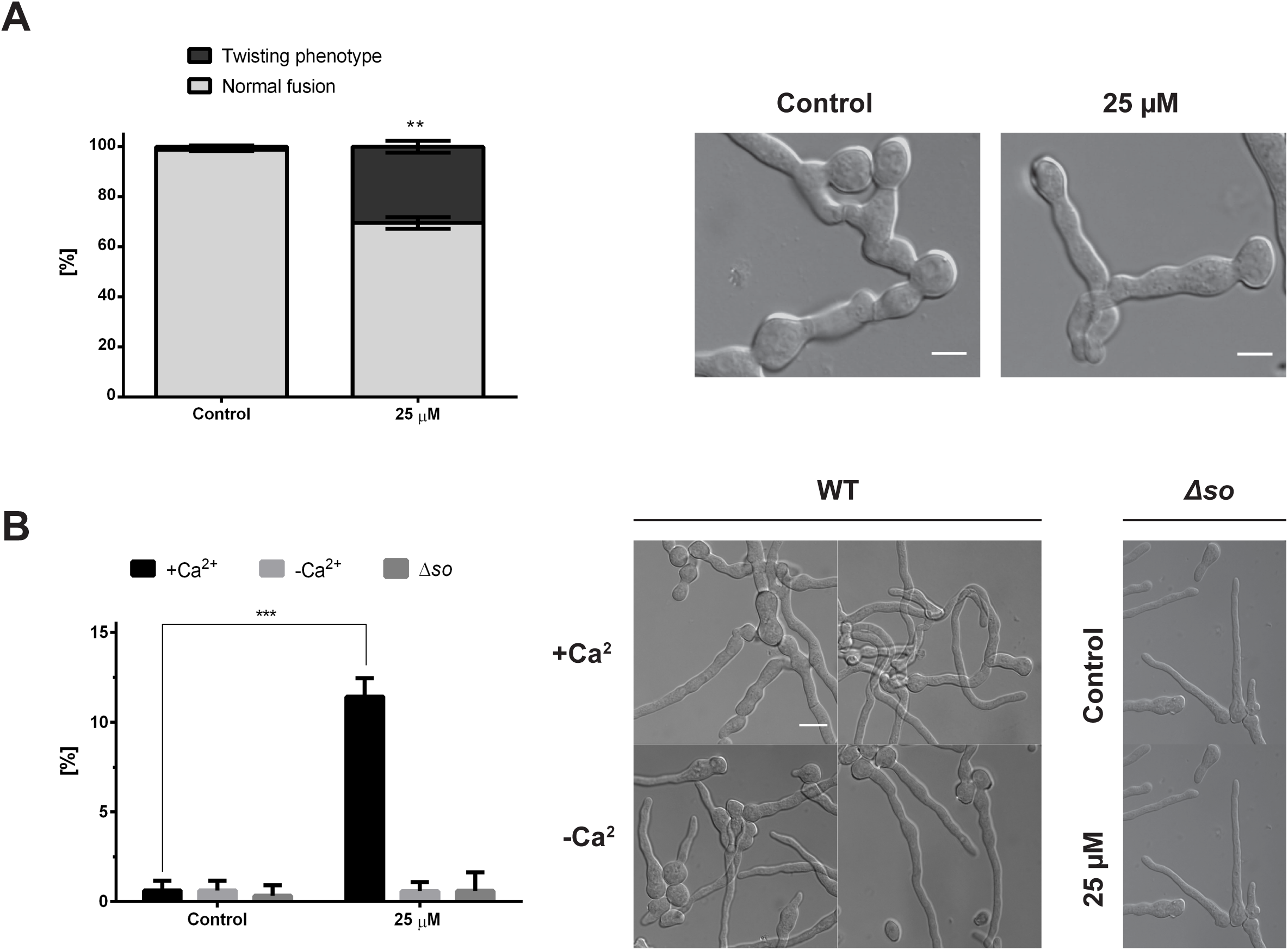
RAC-1 and MAK-1 acts within the same pathway. **(A)** Quantification of twisting and normal growth in cell fusion during partial inhibition of RAC-1. Control (H_2_O) and 25 µM of RAC-1 inhibitor are shown (n≥100). Representative images of both phenotypes are shown at the right. **(B)** Quantification of polarity growth defects when RAC-1 is partially inhibited during cell fusion. Control (H2O) and 25 µM (NSC23766), in media with/without Ca^2+^ and with Δso strain as a control. Significant differences (***, *p*<0.001) were observed between the partial inhibition and the control only in the presence of Ca^2+^.

**Figure 8.**
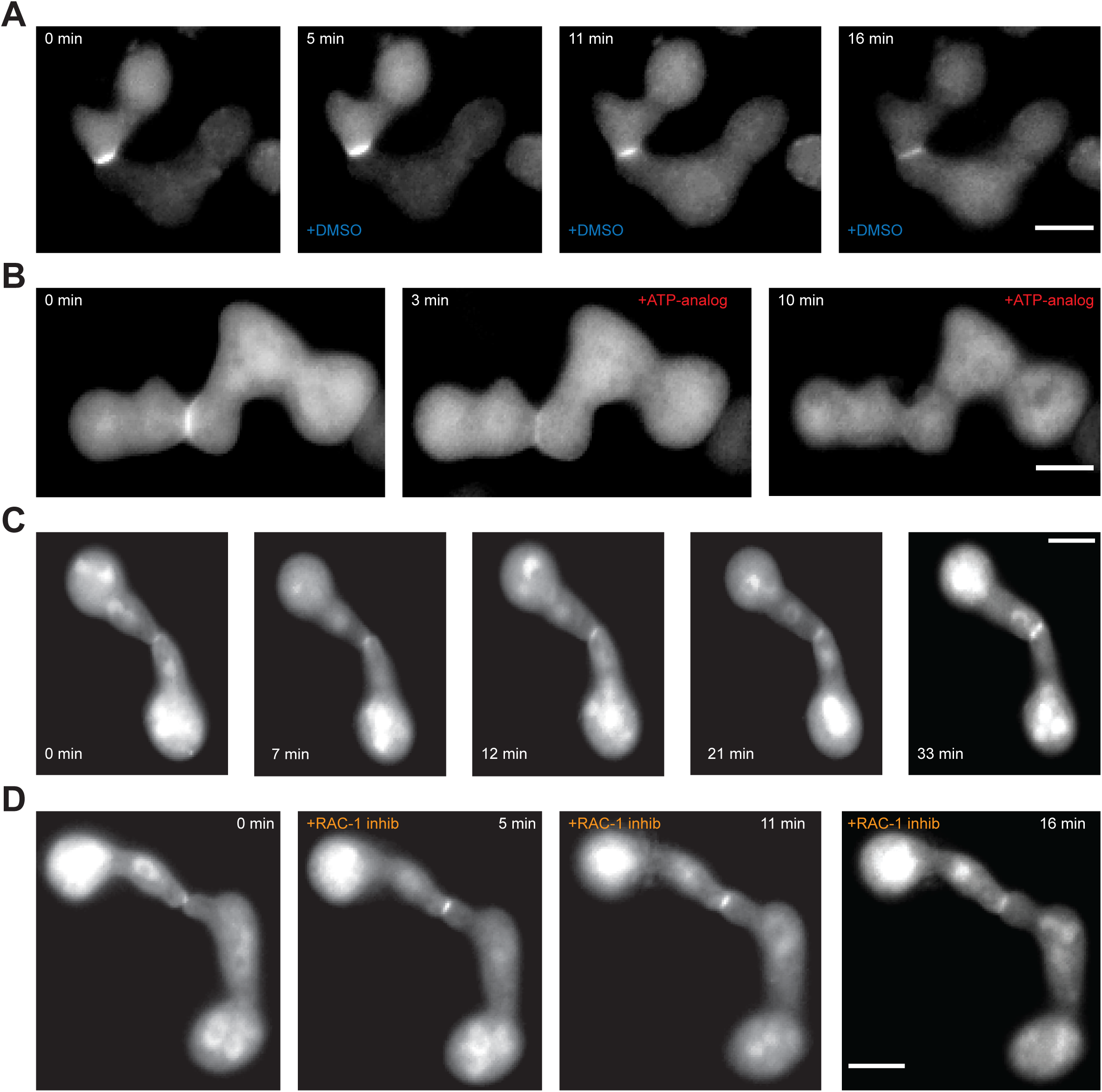
MAK-1 is upstream of RAC-1. **(A)** Localization of CRIB-GFP when cells establish physical contact in a MAK-1^E104G^ background (strain 943), when treated with DMSO. **(B)** Localization of CRIB-GFP during inhibition of MAK-1^E104G^ after cells established physical contact. (*) to point the cell fusion event and signal dissapearence. **(C)** Localization of MAK-1-GFP in a wild-type interaction when cells established physical contact (strain MW513). **(D)** Localization of MAK-1-GFP when RAC-1 is fully inhibited with NSC23766 (100 µM). Similar observations were made multiple times (n=10) Scale bars 5 µm.

These data indicate that the MAPK MAK-1 controls, and is therefore upstream of, the activation and membrane-recruitment of RAC-1 during the cell fusion process.

## Discussion

Cell fusion in filamentous fungi is a minimalist paradigm of morphogenesis and cell communication: genetically identical cells that nevertheless rely on several pathways to convert a shared signal into a directed actin-based growth in an oscillatory manner. In this study we show that the CWI MAPK MAK-1 is essential for activating the Rho GTPase RAC-1 and assembling the actin aster required for chemotropic growth, whereas the MAPK MAK-2 governs the spatial positioning of this RAC-1/Actin cluster. When MAK-1 activity is inhibited, we observed that the actin cluster associated to the tips completely disappeared, arresting the growth of the interacting cells (Fig. 9). Neurospora encodes six Rho-family GTPases, yet only two, RAC-1 and CDC-42, direct polarized growth at the hyphal tip: RAC-1 drives the specialized, chemotropic growth that ends in cell fusion, whereas CDC-42 handles vegetative extension. Our observations match what Lichius *et al*. reported: when RAC-1 is blocked with NSC23766, the RAC-1-active CRIB reporter vanishes, the actin focus dissolves, and growth is arrested, while CDC-42 is unaffected and keeps normal chemotropic growth running [19]. We see the same split, RAC-1 inhibition recreates the MAK-1-block phenotype, but CDC-42 activity and the actin cap at non-interacting tips stay intact, confirming that MAK-1 channels its fusion-specific actin signal through RAC-1 rather than CDC-42. Neurospora carries a single formin, BNI-1, which, like most formins, is normally activated when a GTP-loaded Rho protein binds its regulatory domain, releasing autoinhibition and enabling actin-cable assembly [31]. Surprisingly, however, *bni-1*Δ germlings still undergo fusion and even display a “hyper-fusion” phenotype in which peripheral tips fuse [32]. These observations argue that BNI-1 is not the principal or sole organizer of the fusion actin focus. MAK-1 might therefore influence actin in two, non-exclusive ways: (i) indirectly, by activating RAC-1, which in turn could stimulate BNI-1 activity; or (ii) directly, by phosphorylating BNI-1 (or an alternative actin nucleator) and thereby modulating actin assembly independently of RAC-1. Precedents for both routes exist in higher eukaryotes. The canonical MAPK ERK1/2 phosphorylates the Rho-GEF GEF-H1 to activate the Rho GTPase RhoA [33], while ERK3 not only serves as a GEF for CDC-42/RAC-1 but also phosphorylates the Arp3 subunit of the Arp2/3 complex, boosting actin polymerization. The Arp2/3 complex is the principal catalyst of branched-actin assembly, converting linear filaments into meshworks. In Neurospora, deletion of any Arp2/3 subunit reduces cell-fusion frequency by ∼2.5-fold yet leaves germination unaffected [25], suggesting that Arp2/3, like RAC-1, plays a fusion-specific role and could lie downstream of MAK-1. Arp2/3 may work together with BNI-1 or compensate for it when BNI-1 is absent (in *bni-1*Δ) during fusion. Defining how these nucleators fit into the MAK-1 pathway remains an open question.

**Figure 9.**
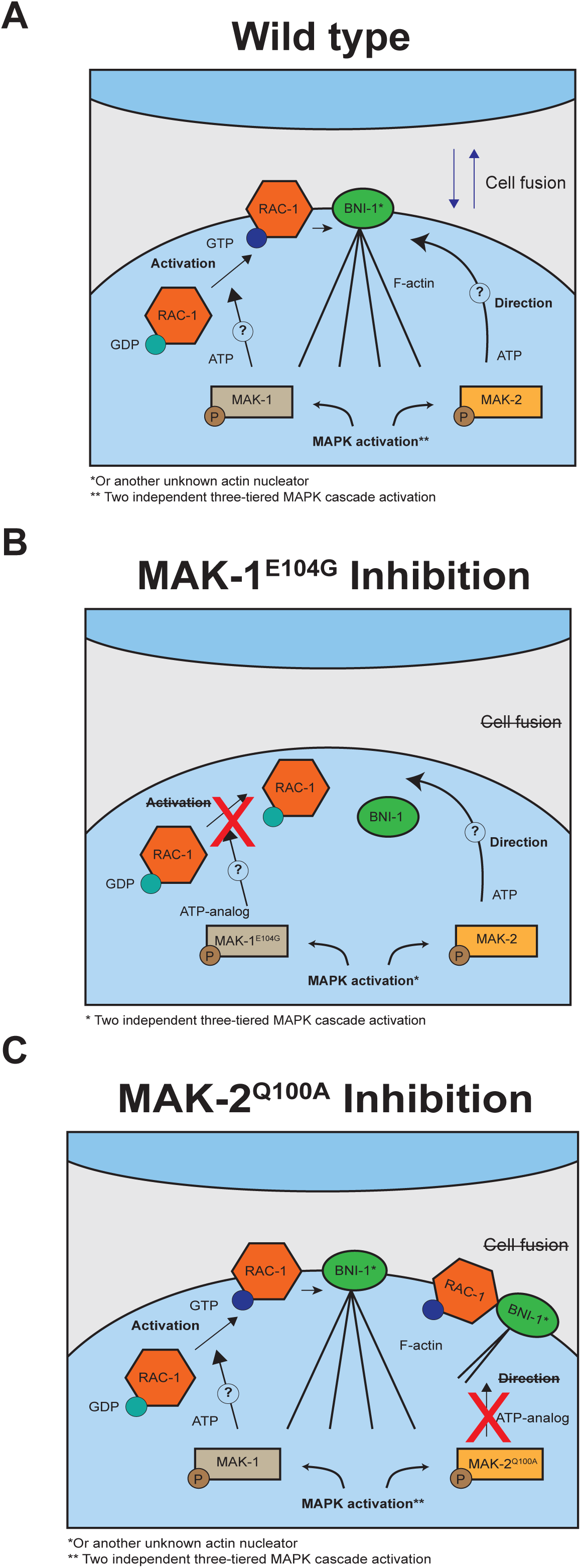
Proposed model of MAK-1/MAK-2 regulation during cell fusion. **A)** During chemotropic growth toward a fusion partner, MAK-1 activates RAC-1, which in turn drives actin polymerization and formation of a focused actin aster at the leading tip. This actin/RAC-1 complex supports polarized exocytosis and directed growth. In parallel, MAK-2 defines the position of this complex by orienting RAC-1 and the actin aster toward the partner cell. **B)** When MAK-1^E104G^ activity is blocked, RAC-1 becomes inactive and detaches from the membrane, the actin aster collapses, and growth stops. **C)** When MAK-2^Q100A^activity is blocked, RAC-1 remains active at the cortex but shifts to ectopic positions around the plasma membrane, and growth loses direction.

Importantly, no evidence currently links CWI MAPKs to direct phosphorylation of RAC proteins. A phosphoproteomic assay of the *Aspergillus fumigatus* CWI MAPK MpkA (the MAK-1 orthologue) catalogued hundreds of MpkA-dependent phosphosites [34]; Supplementary Table S1A) yet did not list the AfRAC-1 (UniProt Q4WWL5) among the modified targets, reinforcing the notion that RAC-type GTPases are controlled indirectly rather than being direct MAPK substrates. Our own data support this view. Inhibiting MAK-1 causes the CRIB-GFP reporter to detach from the plasma membrane, mirroring the loss of RAC-1 signal seen when RAC-1 is chemically blocked with NSC23766 (Fig S6A) [32]. Membrane detachment is diagnostic of RAC-1 inactivation, indicating that MAK-1 activity is required to keep RAC-1 in its GTP-bound state, most plausibly via phosphorylation of an upstream activator. By contrast, inhibiting MAK-2 stops the interaction but leaves RAC-1 on the membrane; the reporter simply relocates to ectopic cortical sites, underscoring MAK-2 role in positioning rather than activating the GTPase. Collectively, these observations favor a model in which MAK-1 sustains RAC-1 activity likely through a yet-to-be-identified GEF or scaffold, while MAK-2 provides the spatial cue that keeps the active RAC-1/actin complex aimed at the fusion partner. A striking parallel comes from the Neurospora ergosterol biosynthesis mutant *erg-2*Δ; perturbing ergosterol composition destabilizes sterol-rich membrane domains (SRDs) at the cell tip and produces the same twisting germ-tube phenotype seen under partial MAK-1 or RAC-1 inhibition (Fig. 7A and B), whereas MAK-2 inhibition simply arrest growth without twisting [20]. SRDs are recognized polarity platforms in several systems: loss of tip-associated SRDs in *Aspergillus fumigatus* or yeast prevent recruitment of polarity organizers and results in growth inhibition [35, 36]. In mammalian cells, Rac1 activation itself requires intact lipid rafts [37]. The matching phenotypes caused by *erg-2*Δ, partial RAC-1 inhibition, and partial MAK-1 inhibition therefore support a model in which SRDs maintain RAC-1 in its active, tip-localised state, most likely via a MAK-1-dependent GEF. Compromising either the SRD scaffold or MAK-1/RAC-1 activity allows RAC-1 to drift from the apex, producing the characteristic twisted morphology seen in *erg-2*Δ germ tubes. This also suggest that inhibition of MAK-2 may scatter the sterol-rich tip patch itself: as the patch drifts around the membrane cortex, it drags the RAC-1/actin complex with it, losing their fixed direction, potentially due to a loss of phase-separation at the cell tips [38], an idea we revisit in the next paragraphs. Previous studies made the hierarchy of the cytoskeleton clear in Neurospora: benomyl-induced loss of microtubules leaves chemotropic interaction unaltered, whereas latrunculin B mediated depolymerization of actin stops tip extension and abolishes fusion [25]. Consistent with this actin dependence, we now see that actin intensity at the two fusion tips oscillates out of phase with the MAK-2/SO signal: actin oscillates independently whether SO or MAK-2 is associated to the tip of the interacting cell. The oscillatory cell dialog has also been described in the nematode-trapping fungus *Arthrobotrys flagrans* [39], but the pattern there is markedly different. In single Arthrobotrys hyphae, in non-interacting cells (described by the authors as a monologue), actin, the SO homologue SofT and the MAK-2 homologue MakB all rise and fall in synchrony, paced by an intrinsic growth oscillator. Only when two hyphae approach do those in-phase oscillations slow, entrain and eventually split into the familiar anti-phasic dialogue between partners [39]. By contrast, *Neurospora* shows no sign of a pre-existing growth oscillator; its actin remains strictly anti-phasic to MAK-2/SO and the rhythm emerges exclusively during cell-to-cell communication. These contrasts imply that the temporal coupling between signaling, and actin assembly has diverged in the two lineages, so insights from Arthrobotrys should be extrapolated to Neurospora with appropriate interpretation.

Mating of the fission yeast *Schizosaccharomyces pombe* offers a well-resolved template for how an actin focus can be integrated with MAPK signaling during fusion. Studies showed that the formin Fus1 nucleates a compact, aster-like actin array, the fusion focus, exactly at the contact site between the two cells [28]. Super-resolution imaging and live secretion assays revealed that this aster acts as a “cargo concentrator,” corralling vesicles laden with cell wall hydrolases so that enzymatic thinning, membrane juxtaposition, and actin-driven force are synchronized. The focus also hosts the pheromone MAPK cascade (MAPK Spk1 and its upstream tiers), creating a signaling hot-spot that locks the cells into the fusion programme. A key mechanistic insight is that the intrinsically disordered region (IDR) of Fus1 undergoes liquid–liquid phase separation: the IDR self-associates to form a condensate whose network simultaneously traps actin regulators, vesicle tethers, and MAPK components in a nanometer-scale volume, greatly boosting their local concentration and reaction rates [38]. Our Neurospora data point to a conceptually similar, though compositionally distinct, hub. The formin BNI-1 is dispensable for the fusion aster, yet two scaffold proteins, HAM-5 and, critically SO, co-localize with the RAC-1/actin complex at the tip. SO is already known to anchor the entire three-tier CWI cascade (including MAK-1) [40], but only SO, not MAK-1, participates in the cell dialog oscillatory dynamic localization, implying that SO fulfils additional, MAPK-independent roles. We therefore propose that MAK-2 (and the signaling cargo it guides) is delivered into a HAM-5 condensate at the apex, where high local concentration of RAC-1 and actin nucleators enforces directionality, in association with SRDs. Whether SO or HAM-5 truly phase-separate, and how MAK-2 activity is choreographed within such a compartment, remains an open question.

Taken together, these oscillation patterns point to species-specific timing circuits that are governed by the upstream MAPKs MAK-1 and MAK-2. Based on our findings we propose a model in which MAK-1 initiates fusion by potentially activating RAC-1 (or perhaps another, yet-unknown actin nucleator) to build the actin aster, whereas MAK-2 continually reorients that aster toward the opposite cell tip (Fig. 9). This system has some similarities with the situation in budding-yeast mating, where the pheromone-response MAPK Fus3 (homolog to MAK-2) redirects growth toward the other partner [41]. Unlike yeast mating, Neurospora somatic cell fusion is pheromone-independent, and its ligand-receptor pair remains unidentified, yet in both systems a dedicated MAPK module converts an external cue into precise spatial guidance of the polarity machinery.

## Material and methods

### N. crassa strains and media

The strains generated and used in this study are listed in the supplementary information (Table S1). Strains were generally grown in Vogel’s Minimal Medium (VMM) (Vogel, 1956) supplemented with 2% sucrose as the carbon source and with 1.5% agar for solid media. Generally, all strains were incubated in solid VMM (supplemented with histidine 0.5 mg/ml when needed) for 3-4 days in dark at 30°C and an additional day with natural light at room temperature. Homokariotic strains were purified by either single spore isolation (SSPi) or by crossing with the wild-type strain. For selection during SSPi, VMM with hygromycin was employed. Crosses were performed on Westergaard’s medium as described earlier (Westergaard and Mitchell, 1947). The ascospores obtained from the crosses were germinated in VMM with hygromycin and confirmed by polymerase chain reaction (PCR).

### Plasmid construction

The primers used for the generation of the plasmids are listed in table S2. In order to construct the *mak-1^E104G^* (analog-sensitive) knock-in cassette, a fragment containing a 5’ upstream region together with the *mak-1* open reading frame was PCR-amplified using the primers 1304 and 1305 and using the previously generated *mak-1^E104G^*-*gfp* plasmid as a template (Weichert et al., 2016). By yeast recombinational cloning, the resulting fragment was fused to the hygromycin resistance cassette (amplified with primers 82 and 83) and a 1 kb fragment homologous to the 3’ downstream region of *mak-1* (generated with primers 1306 and 1307), resulting in the plasmid 738.

For the generation of the *Ptef-1-mak-2^Q100A^* strain, the fragments required for the construction of the gene knock-in cassette were amplified by PCR. A 1 kb genomic region positioned 2 kb upstream of the *mak-2* promoter (considering it to be the 1 kb region upstream of the coding read frame) was amplified with primers 1489 and 1490. The *Ptef-1-mak-2^Q100A^-gfp* plasmid used in a previous study [24] served as a template for the amplification of the *Ptef-1-mak-2^Q100A^* PCR fragment with primers 1488 and 1487. The resistance cassette used was *Hyg^R^*, which was amplified with primers 82 and 83. Finally, the 1 kb genomic region downstream of the *mak-2* Stop-codon was amplified with primers 1489 and 1492. The four fragments were fused by yeast recombinational cloning generating the plasmid 846. The plasmid 723 (*Ptef-1-dsRed-so*) was generated by cloning of the *so* ORF amplified with primers 1419 and 1420 and ligated into the plasmid 722 (pMF334-Tef-dsRed) with enzymes AscI/XbaI. The resulting plasmid was transformed into N1-03 generating the strain 843 (*his-3::Ptef-1-dsRed-so*). Yeast recombinational cloning was performed as previously described (Da et al., 2000).

### Strain generation

All strains were either generated by transformation as described in early studies (Margolin et al., 1997) or by crossing (Westergaard and Mitchell, 1947). When indicated, strains were purchased to the Fungal Genetics Stock Center. A helpful list of protocols and media description is available at their website (www.fgsc.net).

For generating the inhibitable MAK-1 strain, the gatekeeper amino acid, glutamine at position 104, was exchanged with glycine to render the kinase analog-sensitive, resulting in the *mak-1^E104G^* gene [20]. The plasmid 738 containing the *mak-1^E104G^-hph* construct was used as a template to amplify the full-length knock-in fragment by PCR (using primers 1304 and 1307), which was then transformed into the *N. crassa* strain N1-06 (FGSC 9719) (Fig. S1A). This recipient strain has a mutation in the gene *mus52*, which increases the frequency of homologous recombination up to almost 100% [42]. The resulting transformants were first tested by PCR to confirm the base pair exchange (E104G; CAG to GGC), and the proper integration of the transforming DNA was confirmed by Southern blot of homokaryotic isolates obtained from a sexual cross of the primary transformants with the wild-type strain (Fig. S1A and B). To generate the subsequent strains combining reporters and MAK-1^E104G^, strain 849, carrying the *mak-1::mak-1^E104G^-hph* genotype, was crossed independently with strains 665 and 714, which carried the *his-3::Ptef-1-mak-2-gfp* and *his-3::Ptef-1-so-gfp* constructs, respectively, resulting in strains 865 and 874. The integration of the *mak-1::mak-1^E104G^-hph* locus and GFP constructs was confirmed by PCR and fluorescence microscopy, respectively. For LifeAct, strain 849 was crossed with 754 (*his-3::Ptef-1-lifeact-gfp*), resulting in strain 869. The CLA-4 CRIB reporter strain (*Pccg1::crib^cla-4-gfp::bar+*, 927) was crossed with the MAK-1^E104G^ strain (849), resulting in strain 943 (*mak-1::mak-1^E104G^-hph; Pal1-crib^cla-4-bar*). Progenies were tested by fluorescence microscopy and PCR.

Analogous to the design of the *mak-1^E104G^-hph* construct, a *mak-2^Q100A^-hph* gene knock-in cassette was created and integrated at the original gene locus of strain N1-06 (Δ*mus-52*). However, this construct was found to be non-functional, and the respective transformants exhibited a Δ*mak-2*-like phenotype (data not shown). In previous work at our laboratory, we found that the Δ*mak-2* phenotype of the knock-out mutant was only fully complemented when the WT or the analog-sensitive version was overexpressed using strong constitutive promoters such as P*ccg-1* or P*tef-1* [8, 24]. Therefore, the promoter at the original gene locus was replaced by the P*tef-1* sequence. The linear *Ptef-1-mak-2^Q100A^-hph* knock-in cassette, amplified from the plasmid 846 with primers 1487 and 1488, was transformed into N1-05 (Δ*mus-52*, like N1-06 but different mating type), resulting in strain 882 (*Pmak-2-mak-2::Ptef-1-mak-2^Q100A^-hph*). The primary transformants were tested by PCR of the *mak-2* locus and sequencing of the resulting fragment. To obtain a homokaryotic strain, a sexual cross was performed between wild type mat a (N1-02) and one of the primary transformants, 882 (mat A). Four of the resulting progenies were tested by Southern blot analysis for single integration of the construct. All four isolates showed clear bands corresponding to the *Ptef-1-mak-2^Q100A^*radioactive-labelled fragment, while it was absent in the recipient strain (N1-05) (Fig. S4A and B). To generate the subsequent strains combining reporters and MAK-2^Q100A^, we performed a genetic cross between the strain *Pmak-2-mak-2::Ptef-1-mak-2^Q100A^-hph* (899) and two additional strains, 714 (*his-3::Ptef-1-so-gfp*) and 754 (*his-3::Ptef-1-lifeact-gfp*). This generated two new strains: *his-3::Ptef-1-so-gfp; Pmak-2-mak-2::Ptef-1-mak-2^Q100A^-hph* (912) and *his-3::Ptef-1-lifeact-gfp; Pmak-2-mak-2::Ptef-1-mak-2^Q100A^-hph* (915), respectively. The CRIB reporter strain was generated by crosses of 927 (*Pal1-crib^cla-4-bar*) with MAK-2^Q100A^; 899 (*Pmak-2-mak-2::Ptef-1-mak-2^Q100^A-hph*), resulting in strain 949 (*Pmak-2-mak-2::Ptef-1-mak-2^Q100^A-hph;Pal1-crib^cla-4-bar*).

An actin-labeled Δ*mak-1* strain was obtained by setting up a sexual cross between the Δ*mak-1* (NCU09482; purchased to the Fungal Genetics Stock Center; www.fgsc.net) and 754 (*his-3::Ptef-1-lifeact-gfp*), resulting in the 891 strain (*mak-1::hph;his-3::Ptef-1-lifeact-gfp*).

### Live-cell imaging

Sample preparation was performed as previously described (Schürg et al., 2012). For observation of hyphal fusion in *N. crassa*, 5 μl of a 10^7^-spore suspension were inoculated on the side of a MM agar plate and incubated for 15 hours at 30°C. The preparation of the sample was performed in a similar way than for the spores. The microscope used and the settings for software analyses were described in an early study (Serrano et al., 2018). All fluorescence images shown in this study were processed by the widely used commercial deconvolution software Huygens (by Scientific Volume Imaging), more specifically the Huygens Essential version. Z-stacks (usually n=10) up to 100 nm of all fluorescence images were captured and assembled as a single tiff file by using ImageJ (image-stacks-image to stack). The composite images were processed through the deconvolution software with the following parameters (40-100 iterations, 12 signal/noise ratio and 0.01% of quality change thresh).

### Quantitative analyses

Quantification of the cell fusion and germination rate was performed as described elsewhere (Fleissner et al., 2009; Schürg et al., 2012). For all cell fusion quantifications at least 100 spores were quantified per sample/condition, and at least three independent repetitions were studied.

### Chemical inhibition

Three different chemicals were used for the corresponding experiments. Latrunculin A (10 µM), dissolved in water, was used to test the consequences of actin-polymerization inhibition, 1-NM-PP1 (40 µm) dissolved in DMSO was used to test the inhibition of the analog-sensitive kinases, and NSC23766 (100 µM/25 µM), dissolved in water, was used for specific inhibition of the Rho-GTPase RAC-1. As a control, either water or DMSO 1% was used. For all inhibitor experiments, fresh spores were harvested and incubated for 2 hours at 30°C in MM plates. A 1 cm^2^ piece was cut from the agar plate and inverted onto a coverslip. The chemical was added on the sample prior inversion onto the coverslip or on one side of the agar block.

### Statistics applied to the data

All quantitative data were statistically analyzed by performing a two-tailed unpaired t-test using Excel. The P values were calculated for 0.001, 0.05 or 0.01, and indicated as ***, ** or * respectively. To analyze whether the variances were paired or unpaired, F-test analyses were performed.

## Supplementary information

### Supplementary figures

**Figure S1. Characterization of MAK-1^E104G^ strains. (A)** Graphical representation of the knock-in construct integrated in the genome of the transformant strains. Arrows indicate BamH1 cutting sites and the bands that are generated. **(B)** Southern blot of the positive strains obtained after transformation and crossing with the wild-type strain. DNA was treated with BamH1 before hybridization with the membrane **(C)** Macroscopic phenotype of wild type and MAK-1^E104G^ (strain 849) in presence of DMSO or ATP-analog. Similar observations were made on strains 848, 850 and 851 (same genotype than 849). **(D)** Quantification of tropic interactions and germination of MAK-1^E104G^ strain in presence of DMSO or the ATP-analog (n>100).

**Figure S2. Oscillatory recruitment of MAK-2/SO and actin is disrupted after MAK-1 inhibition during hyphal fusion. (A)** Localization of MAK-2-GFP during inhibition of MAK-1^E104G^ (strain 865) in a hyphal fusion. **(B)** Localization of SO-GFP during inhibition of MAK-1^E104G^ (strain 874) in a hyphal fusion. **(C)** Localization of LifeAct-GFP during inhibition of MAK-1^E104G^ (strain 869) in a hyphal fusion. Similar observations were made multiple times (n>5). Scale bars: 5 µm.

**Figure S3. Quantifications of signal intensity during cell fusion. (A)** Lifeact relative intensity ratio between the signal localized at 3 µm within the apex of the cell and the cytoplasm, from the images shown in figure 2A. **(B)** Lifeact relative intensity ratio between the signal localized at 3 µm within the apex of the cell and the cytoplasm, from the images shown in figure 2B. **(C)** SO relative intensity ratio between the signal localized at 3 µm within the apex of the cell and the cytoplasm, from the images shown in figure 2B. **(D)** Combination of data extracted from graphs B and C from the left cell of the image, indicating a asynchronous oscillation.

**Figure S4. Characterization of MAK-2^Q100A^ strains. (A)** Graphical representation of the knock-in construct integrated in the genome of the transformant strains. Arrows indicate BamH1 cutting sites and the bands that are generated. **(B)** Southern blot of the positive strains obtained after transformation and crossing with the wild-type strain. DNA was treated with BamH1 before hybridization with the membrane. **(C)** Macroscopic phenotype of wild type and MAK-2^Q100A^ (strain 898) in presence of DMSO or the ATP-analog. Similar observations were made on strains AS899, AS900 and AS901 (same genotype as 898). **(D)** Quantification of tropic interactions and germination of MAK-2^Q100A^ strain in presence of DMSO or the ATP-analog (n>100).

**Figure S5. Actin and MAK-2 inhibition effect on SO/actin dynamics. (A)** Localization of SO-GFP during MAK-2^Q100A^ (strain 912) inhibition on interacting cells. **(B)** Localization of LifeAct-GFP during MAK-2^Q100A^ (strain 915) inhibition on interacting cells. **(C)** Effect of the actin inhibitor latrunculin A on actin assembly in LifeAct-GFP localization (strain 754). **(D)** Effect of RAC-1 inhibition on LifeAct-GFP (strain 869) localization during cell fusion. Similar observations were made multiple times (n>5).

**Figure S6. Effect of RAC-1 inhibition on CRIB-GFP localization and actin assembly on strain. (A)** CRIB-GFP localization after RAC-1 inhibition (NSC23766) during cell fusion. **(B)** Relative intensity ratio between the signal localized at 3 µm within the apex or in the germ tube of the cell and the cytoplasm, from the images shown in figure 5C. At the left *y-axis* is represented the apex/cytop signal (red and dark blue) and in the right *y-axis* blue) and in the right *y-axis* is represented the germ tube plasma membrane/cytop signal (light blue and orange). (C) Actin localization and organization in Δ*mak-1* (strain NCU09482).

**Table S1.**
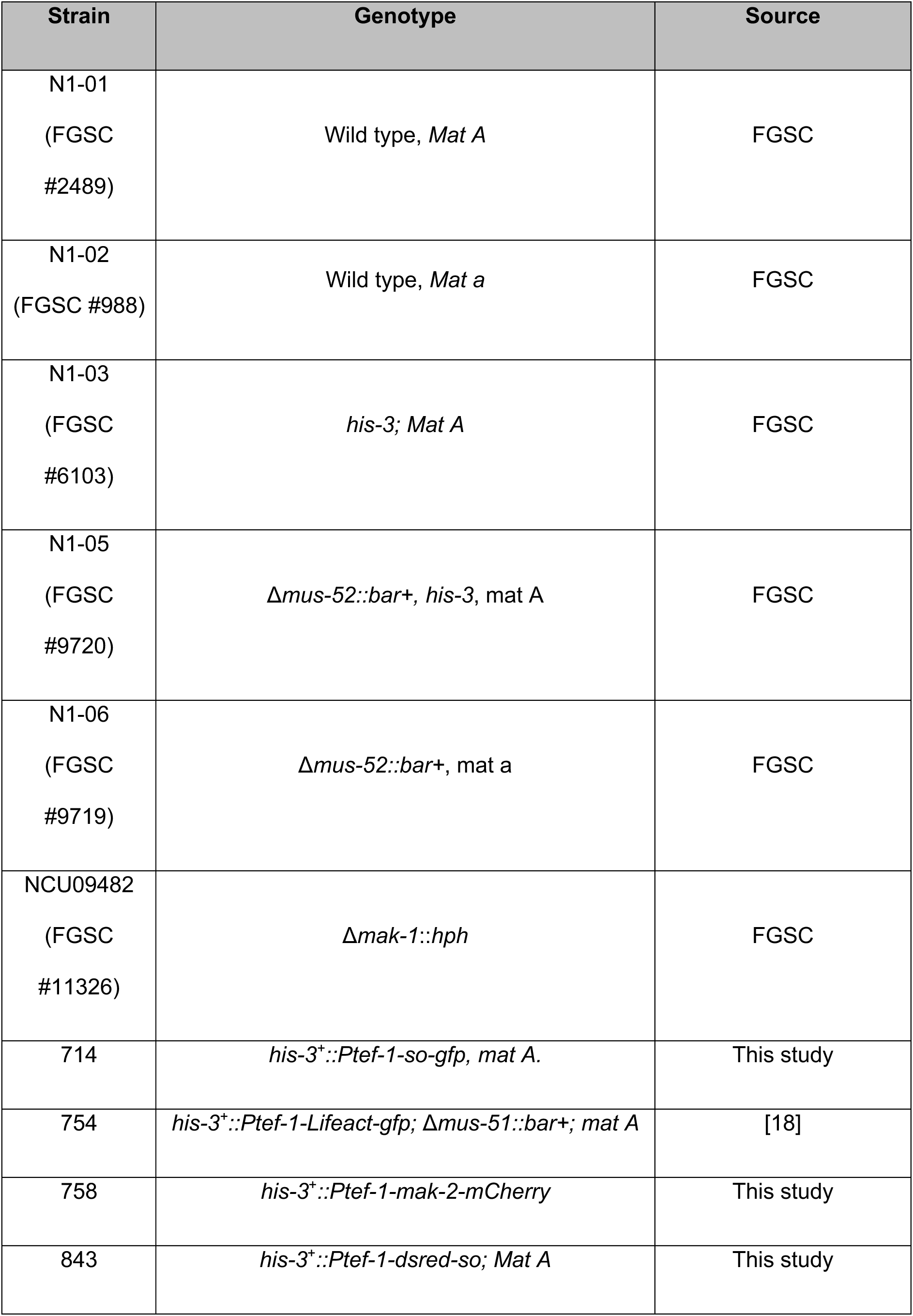
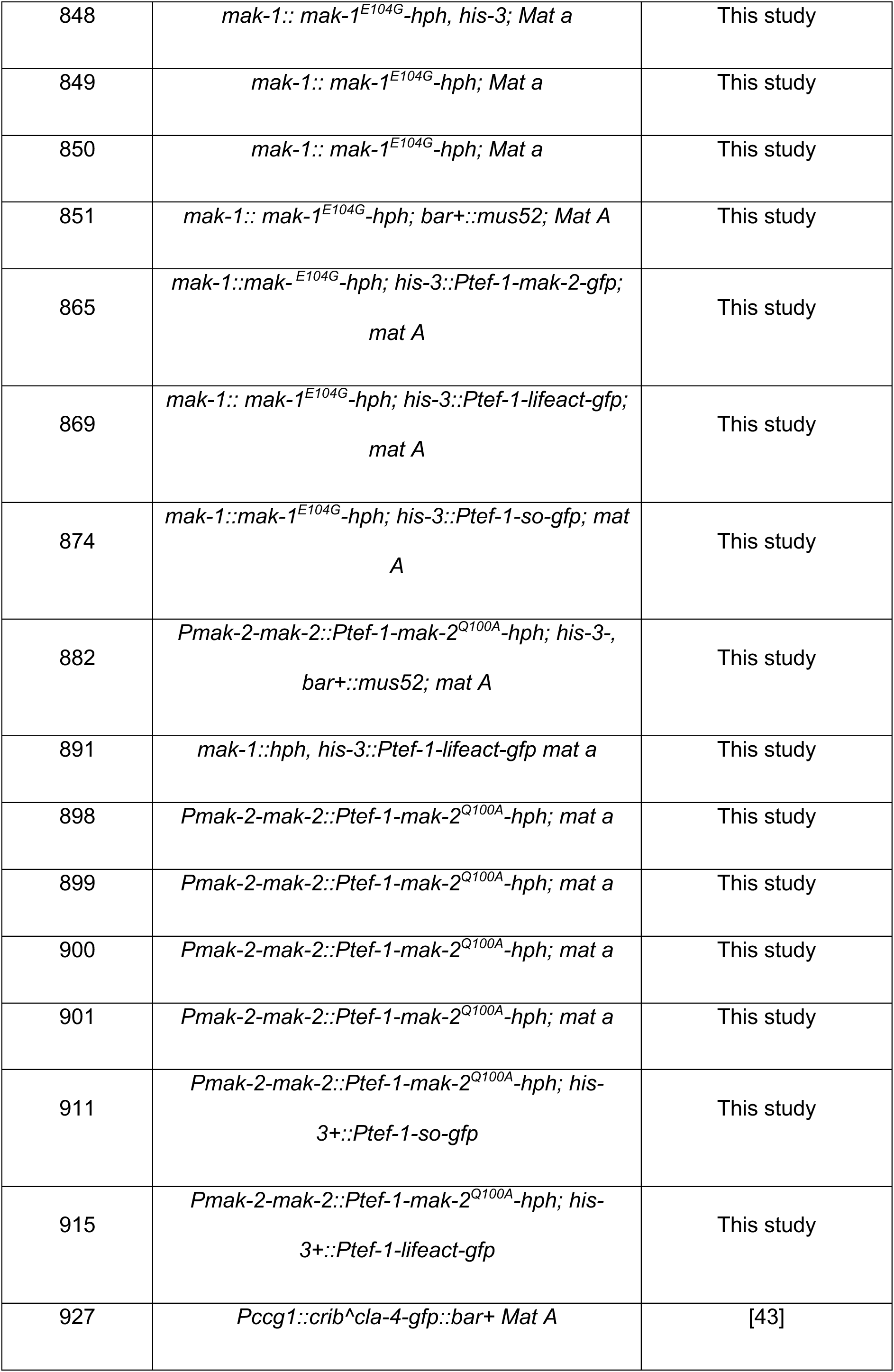
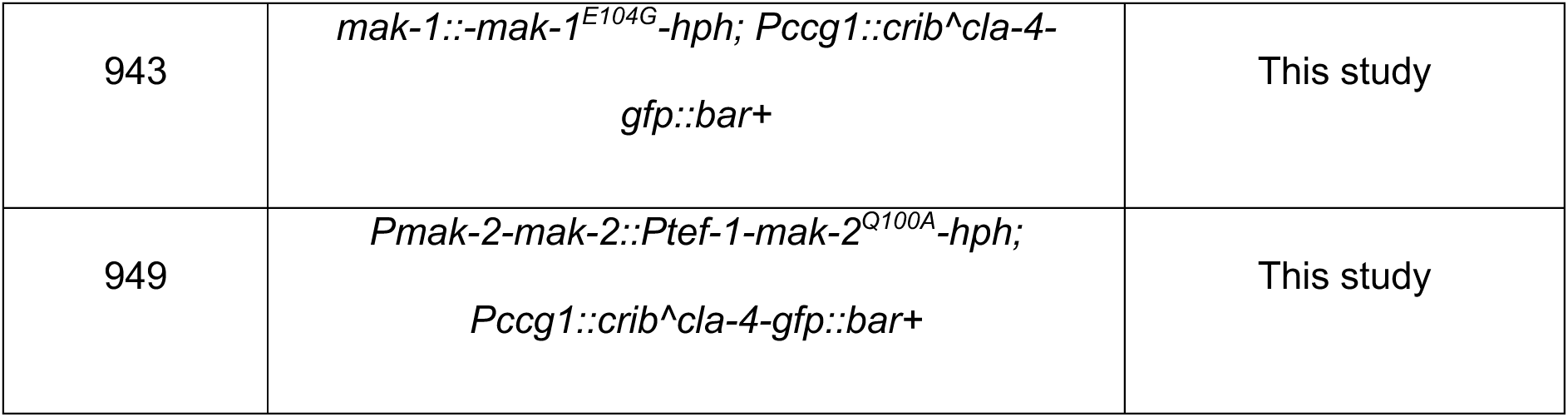
List of strains used in this study.

**Table S2.**
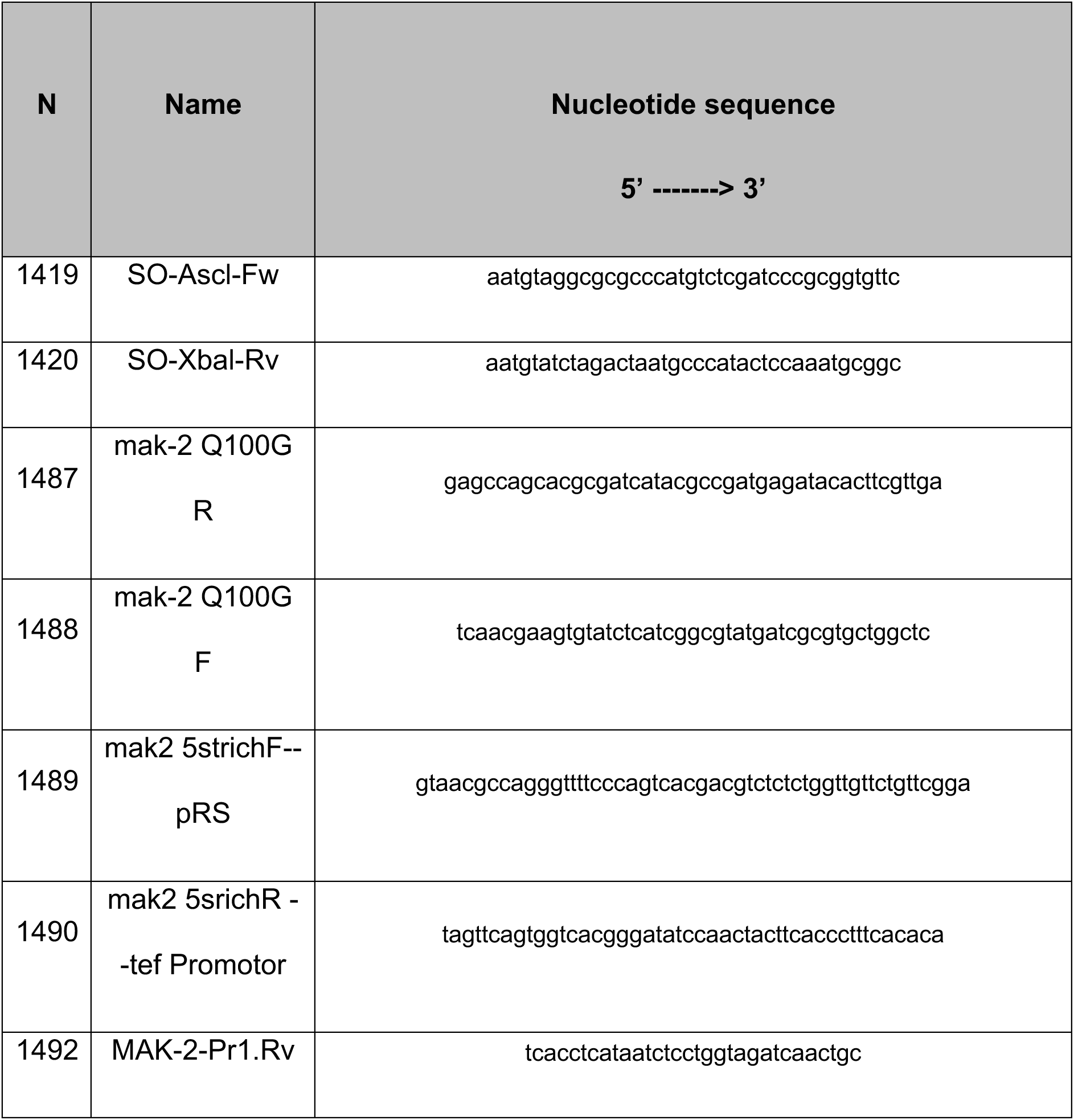
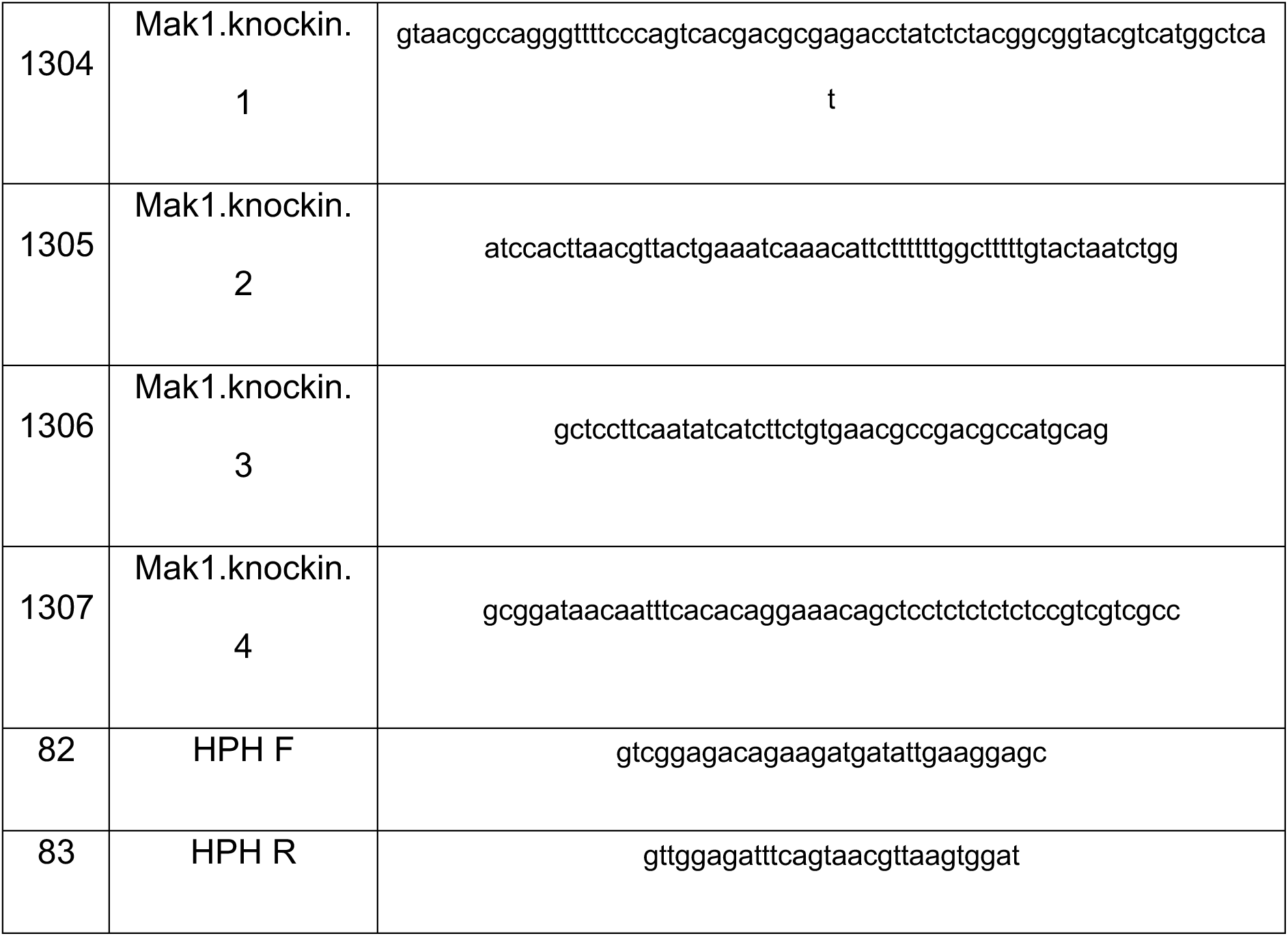
List of primers used in this study.

**Table S3.**
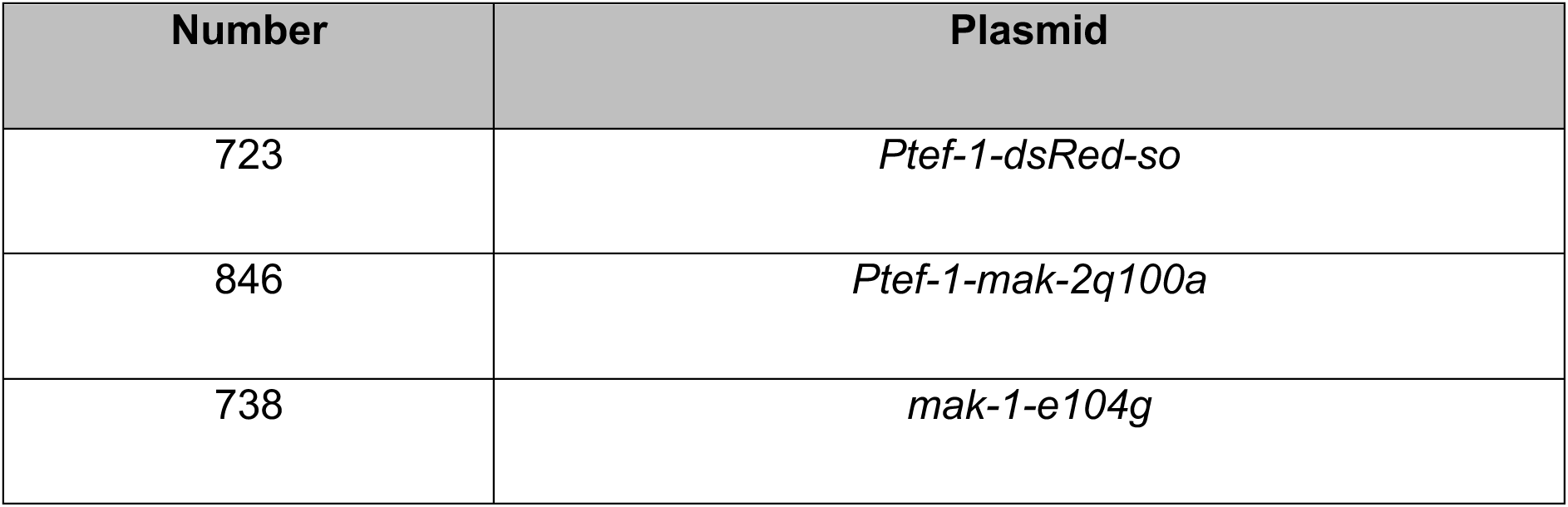
List of plasmids used in this study.

| Number | Plasmid |
| --- | --- |
| 723 | *Ptef-1-dsRed-so* |
| 846 | *Ptef-1-mak-2q100a* |
| 738 | *mak-1-e104g* |

## Notes

### Competing Interest Statement

The authors have declared no competing interest.

### Summary of Updates

Full manuscript updated and revised. Introduction, results and discussion revised. New figure added (figure 9) including a proposed model of the MAK-2 & MAK-1 roles in somatic cell fusion.

