## Supplementary figures and images for "Directed growth during somatic cell fusion in *Neurospora crassa* requires contributions from two distinct MAP kinase pathways"

### Supplementary figures

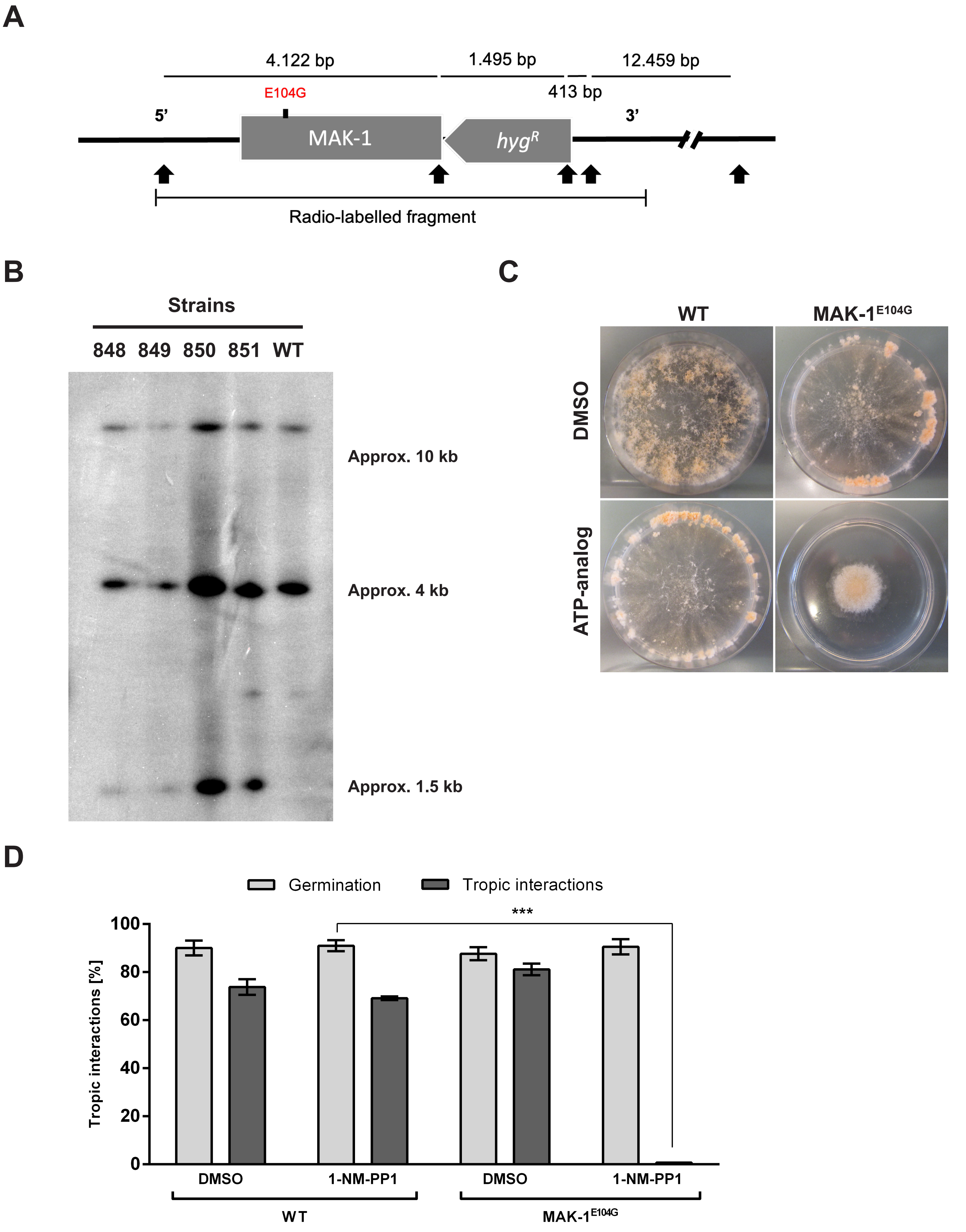

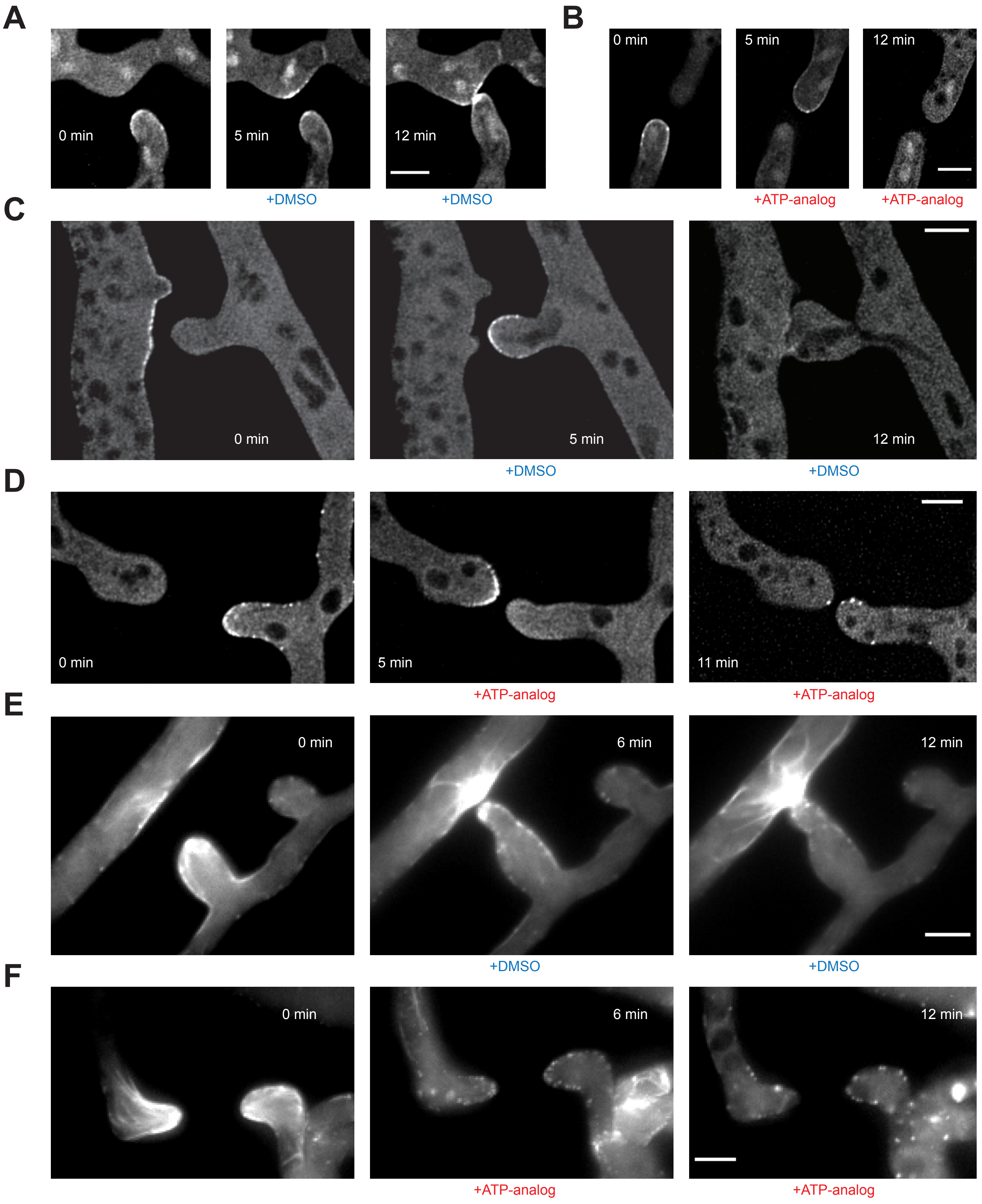

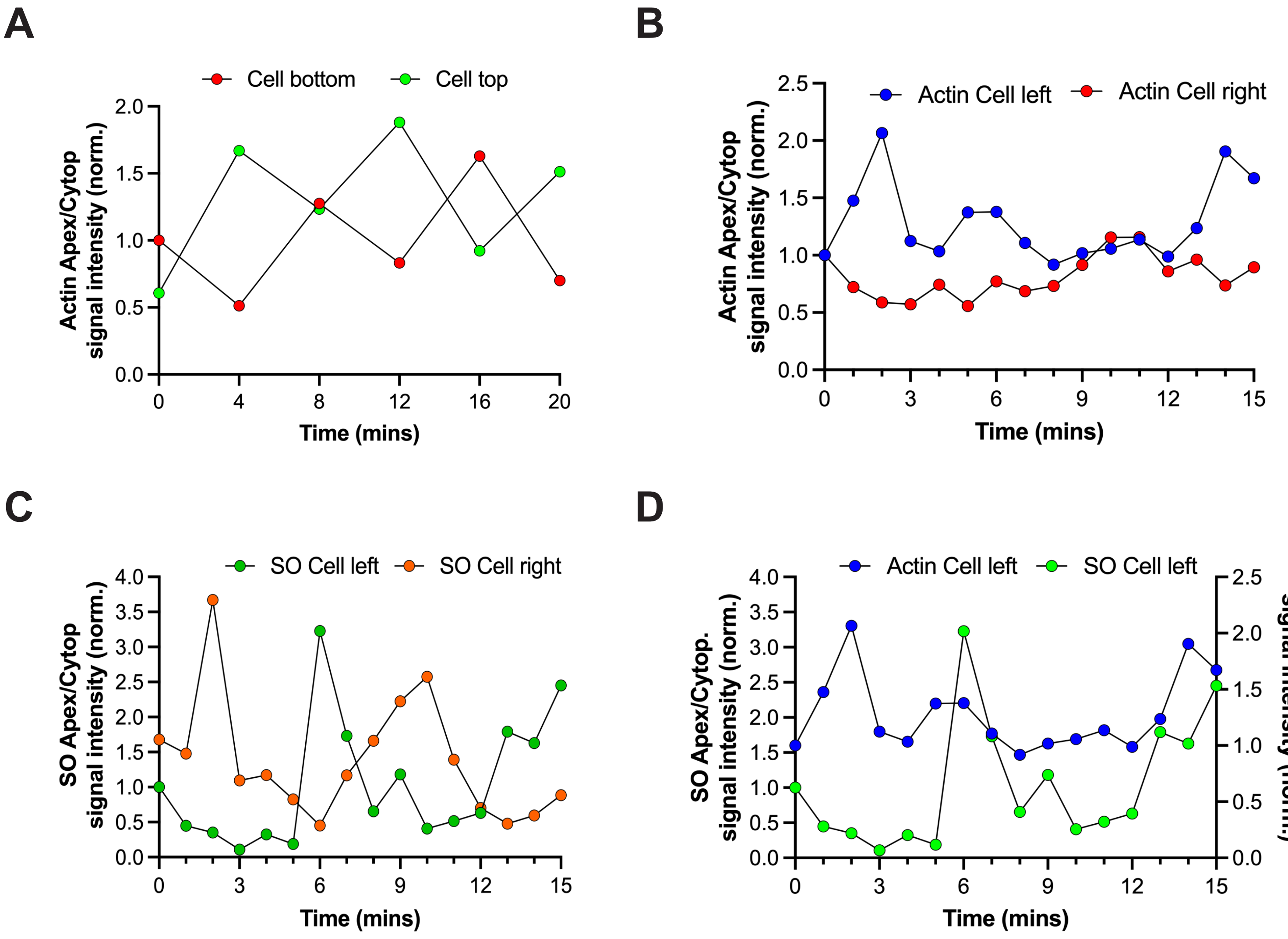

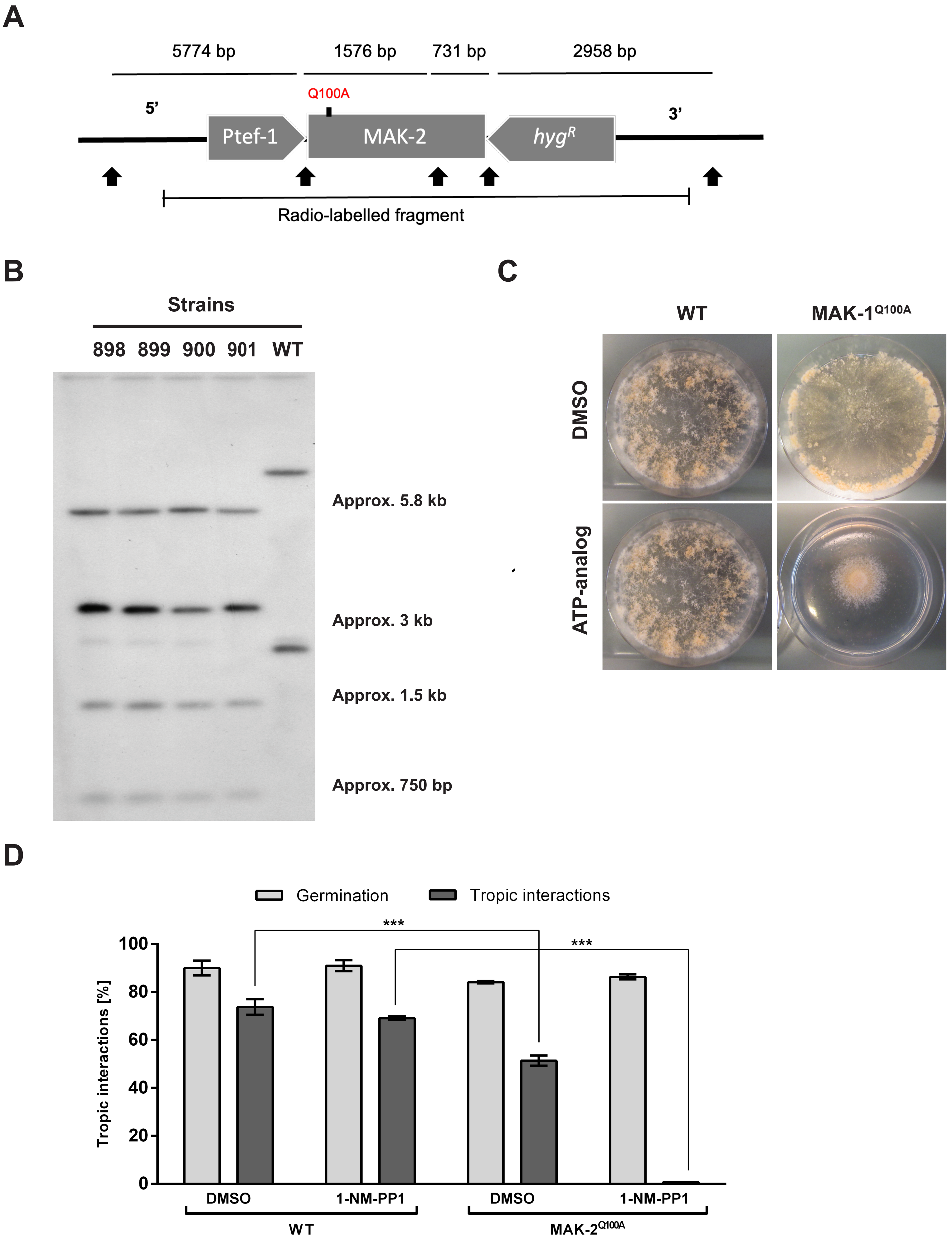

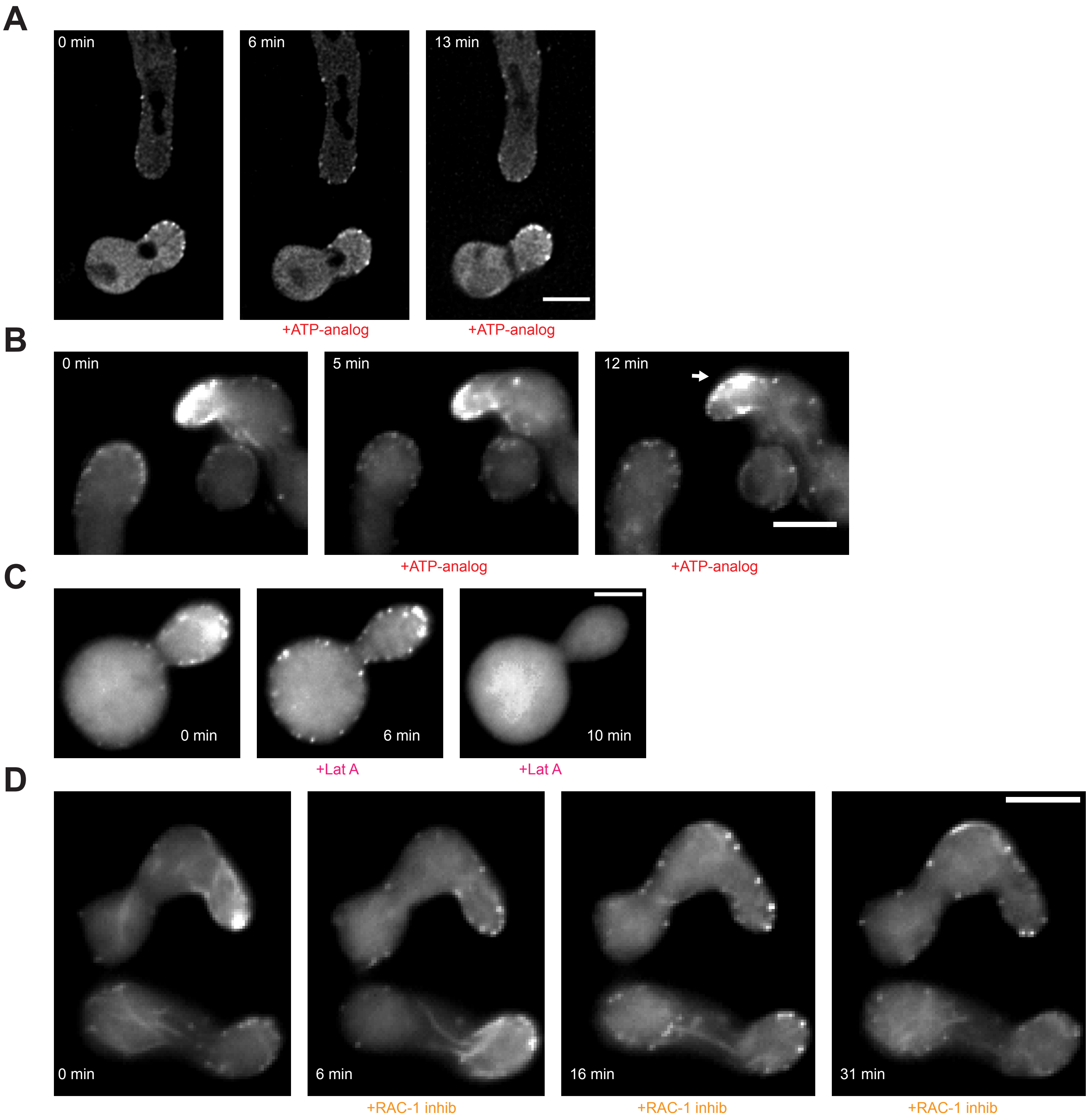

A

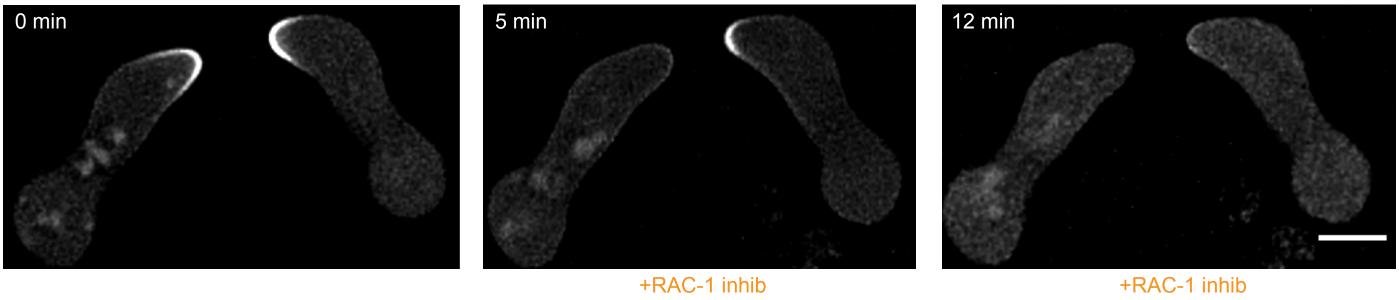


B

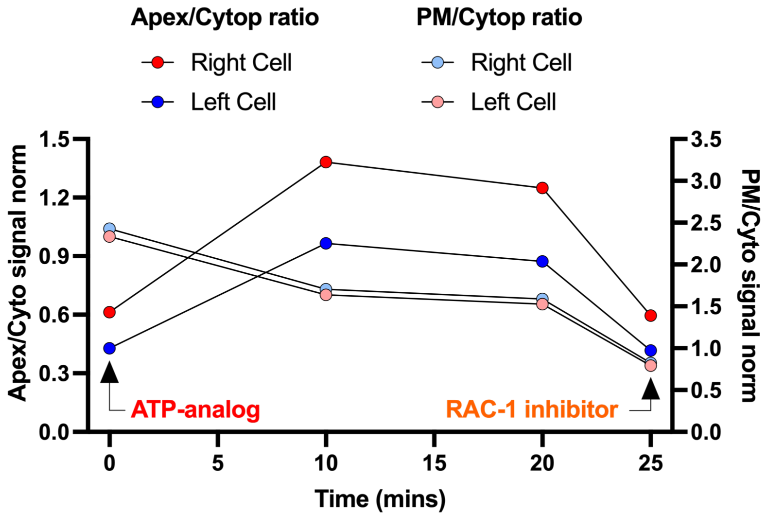


C

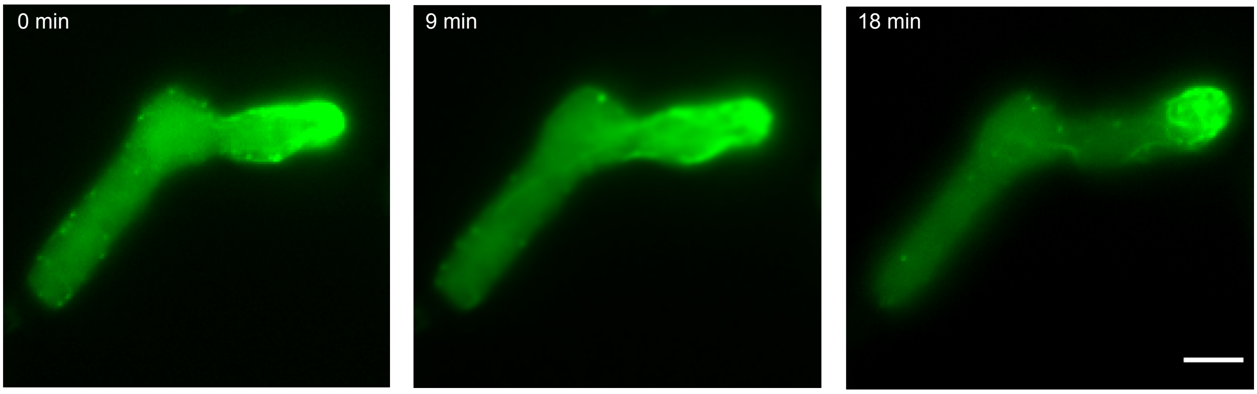


#### Figure S6
